# A synthetic synaptic organizer protein restores glutamatergic neuronal circuits

**DOI:** 10.1101/2020.02.27.967836

**Authors:** Kunimichi Suzuki, Jonathan Elegheert, Inseon Song, Hiroyuki Sasakura, Oleg Senkov, Wataru Kakegawa, Amber J. Clayton, Veronica T. Chang, Maura Ferrer-Ferrer, Eriko Miura, Rahul Kaushik, Masashi Ikeno, Yuki Morioka, Yuka Takeuchi, Tatsuya Shimada, Shintaro Otsuka, Stoyan Stoyanov, Masahiko Watanabe, Kosei Takeuchi, Alexander Dityatev, A. Radu Aricescu, Michisuke Yuzaki

**Affiliations:** Department of Physiology, Keio University School of Medicine, Tokyo 160-8582, Japan; Division of Structural Biology, University of Oxford, Oxford OX3 7BN, UK; Molecular Neuroplasticity, German Center for Neurodegenerative Diseases (DZNE), 39120 Magdeburg, Germany; Department of Medical Cell Biology, School of Medicine, Aichi Medical University, Aichi, Japan; Neurobiology Division, MRC Laboratory of Molecular Biology, Cambridge CB2 0QH, UK; Medical Faculty, Otto-von-Guericke-University, 39120 Magdeburg, Germany; Center for Behavioral Brain Sciences (CBBS), 39106 Magdeburg, Germany; Department of Anatomy, Hokkaido University Graduate School of Medicine, Sapporo 060-8638, Japan

## Abstract

Neuronal synapses undergo structural and functional changes throughout life, essential for nervous system physiology. However, these changes may also perturb the excitatory/inhibitory neurotransmission balance and trigger neuropsychiatric and neurological disorders. Molecular tools to restore this balance are highly desirable. Here, we report the design and characterization of CPTX, a synthetic synaptic organizer combining structural elements from cerebellin-1 and neuronal pentraxin-1 to interact with presynaptic neurexins and postsynaptic AMPA-type ionotropic glutamate receptors. CPTX induced the formation of excitatory synapses *in vitro* and *in vivo* and restored synaptic functions, motor coordination, spatial and contextual memories, and locomotion in mouse models for cerebellar ataxia, Alzheimer’s disease and spinal cord injury, respectively. Thus, CPTX represents a prototype for novel structure-guided biologics that can efficiently repair or remodel neuronal circuits.

**One Sentence Summary:** Structural biology information was used to design CPTX, a synthetic protein that induces functional excitatory synapses and restores normal behaviors in mouse models of neurological diseases.

## Main Text

A broad range of neuropsychiatric and neurological disorders, including autism spectrum disorders, epilepsy, schizophrenia and Alzheimer’s disease, are thought to be caused by an imbalance between excitatory and inhibitory (E/I) synaptic functions (*1–5*). During organism development, but also throughout life, the formation and remodeling of complex yet precise neuronal circuits rely on specific synaptic organizing proteins (*6, 7*). These include cell adhesion molecules, such as neurexins (Nrxs), neuroligins (*8, 9*) and receptor protein tyrosine phosphatases (*10*), as well as secreted proteins, such as fibroblast growth factors, semaphorins, Wnt and extracellular scaffolding proteins (ESPs) (*7, 11, 12*). ESPs directly connect pre- and postsynaptic membrane proteins to form molecular bridges that span the synaptic cleft and mediate bi-directional signaling. For example, cerebellin-1 (Cbln1) is released from parallel fibers (PFs; axons of cerebellar granule cells) and contributes to the synapse-spanning tripartite complex Nrx/Cbln1/GluD2 (the ionotropic glutamate receptor family member delta-2) (Fig. 1A, left) by simultaneously binding Nrx isoforms containing the 30-residue “spliced sequence 4” (SS4) insert (Nrx(+4)) expressed at PF terminals and the amino-terminal domains (ATD, also known as NTD) of GluD2 on Purkinje cells (PCs) (*13–15*). Importantly, a single injection of recombinant Cbln1 into the cerebellum can restore ∼75% PF–PC synapses and normal motor coordination within 1 d after injection in adult *Cbln1*-null mice *in vivo* (*16*). Cerebellins (Cbln1–4) are expressed in nearly all brain regions with distinct patterns and developmental dynamics (*17, 18*). Cbln1 promotes the formation or maintenance of excitatory synapses in the nucleus accumbens (*19*), while Cbln2 is necessary for excitatory synapses in the interpeduncular nucleus (*20*) and the hippocampus (*21, 22*). Cbln4 mediates formation of inhibitory synapses between pyramidal neurons and cortical interneurons (*23*) by interacting post-synaptically with GluD1 (*24*), a receptor closely related to GluD2 (*25*). The C1q-related ESPs C1ql1 promotes excitatory synapse formation and maintenance in the cerebellum by binding to the adhesion G-protein coupled receptor BAI3 (*26, 27*). C1ql3 also mediates synapse formation and maintenance in the medial prefrontal cortex (*28*), while C1ql2 and C1ql3 produced by mossy fibers recruit kainate-subtype ionotropic glutamate receptors (KAR) by directly binding to the ATDs of GluK2/4 KAR subunits in CA3 hippocampal neurons (*29, 30*). Molecular components of excitatory synapses are considerably different among neuronal circuits (*31, 32*) and it is likely that other ESPs remain to be discovered. Nevertheless, the Cbln1–Cbln4 and C1ql1–C1ql3 examples prompted us to hypothesize that synthetic molecules with pre-determined pre- and post-synaptic binding specificities could be designed and employed to reverse the loss of synapses and promote the structural and functional recovery of damaged neuronal circuits.

**Fig. 1.**
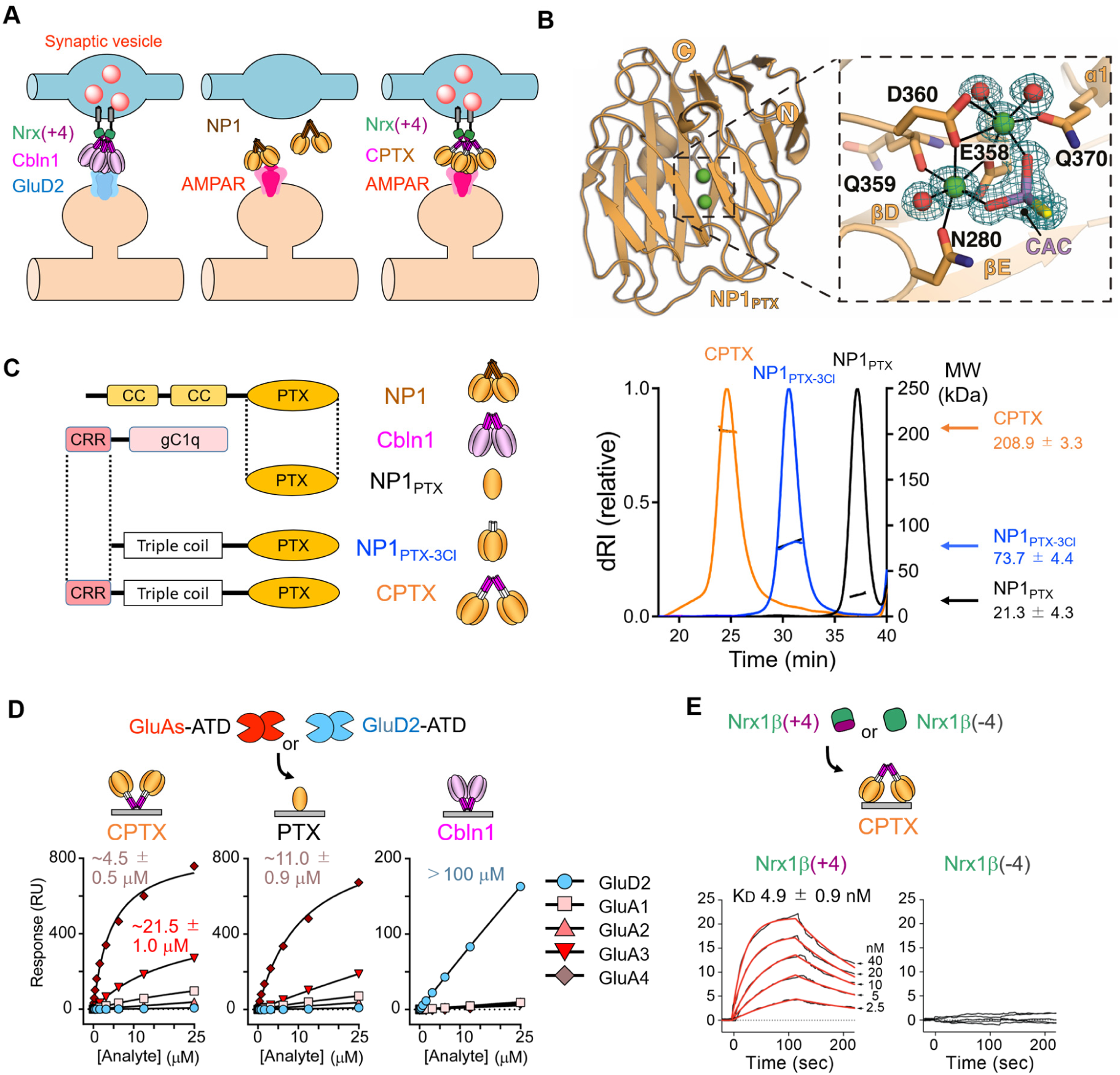
Design of CPTX guided by the structures of NP1 pentraxin domain and Cbln1. **(A)** The CPTX concept. While Cbln1 induces synapse formation by binding to postsynaptic GluD2 and presynaptic Nrx(+4), NP1 induces clustering of postsynaptic AMPARs without inducing presynaptic elements. CPTX was designed as a chimeric ESP of Cbln1 and NP1. **(B)** Crystal structure of the pentraxin domain of human NP1. The N- and C-termini are annotated. Calcium (Ca) atoms and water molecules are shown as green and red spheres, respectively. 2mFo-DFc electron density is contoured at 1.0 σ. The Ca^2+^ coordination shell, which includes a cacodylate buffer molecule from the crystallization solution, is indicated by black lines. **(C)** Diagram of CPTX and related constructs. CC, coiled-coil domain; PTX, pentraxin domain, CRR, cysteine-rich region; gC1q, global C1q domain; 3Cl, triple coil. The right graph shows molecular masses of monomeric NP1_PTX_, trimeric NP1_PTX-3Cl_ and hexameric CPTX. **(D)** Binding isotherms for the interactions between GluA1–4 ATD or GluD2 ATD and immobilized CPTX, NP1_PTX_ and Cbln1. The ATD of GluA4 interacted strongest with CPTX (*K*_D_ ∼ 4.5 ± 0.5 μM) and NP1_PTX_ (*K*_D_ ∼ 21.5 ± 1.0 μM), but not with Cbln1. In contrast, the ATD of GluD2 only interacted with Cbln1 (*K*_D_ >100 μM). **(E)** Single-cycle kinetic sensorgrams for the interaction between the ectodomain of Nrx1β(±4) and immobilized CPTX. CPTX directly bound to Nrx1β(+4) (*K*_D_ ∼ 4.90 ± 0.90 nM), but not to Nrx1β(–4).

### Design of a synthetic ESP-type synaptic organizer

Neuronal pentraxins (NPs) are a family of oligomeric secreted (NP1 and NP2, also known as NARP) or membrane-attached (NPR) proteins that induce clustering of postsynaptic AMPA-type ionotropic glutamate receptors (AMPARs) through direct interactions between pentraxin (PTX) domains and the AMPAR ATDs (*33–36*). Astrocyte-secreted glypican 4 indirectly leads to AMPAR recruitment by enhancing the release of NP1 from pre-synaptic axons (*37*). However, unlike Cbln1 (*13, 16*) and Cbln4 (*23, 24*), there is no evidence indicating that NPs induce presynaptic specializations *in vivo* (*38, 39*) (Fig. 1A, middle). Thus, we hypothesized that we could exploit the modular architecture of NP1 and Cbln1 to develop a synthetic hexameric ESP, termed CPTX, which would include the N-terminal cysteine-rich region (CRR) of Cbln1 that binds Nrx(+4) and the PTX domain of NP1 (Fig. 1A, right). CPTX should have the potential to induce the formation of excitatory synapses and may organize trans-synaptic nanocolumns (*40*) by directly recruiting presynaptic vesicle release machinery (though Nrx binding) and postsynaptic AMPARs (through PTX binding).

To define the correct domain boundaries for CPTX assembly, we first solved the crystal structure of the PTX domain of NP1 (NP1_PTX_) at 1.45 Å resolution (Fig. 1B and Table S1). Reminiscent of short pentraxins (*41*), such as the C-reactive protein and serum amyloid P component, NP1_PTX_ formed a two-layered β sheet with a flattened jellyroll topology containing two Ca^2+^ ions (Fig. 1B and fig. S1). However, unlike short pentraxins, NP1_PTX_ is monomeric in the crystal as well as in solution, as confirmed by multi-angle light scattering (MALS; Fig. 1C). Surface plasmon resonance (SPR) assays, performed to identify the minimal domain requirements for NP1-GluA interactions, revealed that NP1_PTX_ bound the ATDs of GluA1, GluA3 and GluA4 AMPARs with affinities in the high μM range (Fig. 1D). Our recent analysis of the Nrx/Cbln1/GluD2 trans-synaptic complex showed a similarly weak interaction between the globular domain of Cbln1 and the GluD2 ATD; however, avidity effects arising from the oligomeric nature of Cbln1 (hexamer) and GluD2 (tetramer) increase the apparent affinity between the full-length partners considerably (*15*). To mimic the structural organization of Cbln1 in CPTX, we first added a triple-coil-forming mutant GCN4 peptide to the amino-terminus of NP1_PTX_, resulting in the NP1_PTX-3Cl_ construct, and confirmed its trimeric stoichiometry (Fig. 1C). The Cbln1 CRR (*15*) was then added to the NP1_PTX-3Cl_ N-terminus, leading to the hexameric CPTX molecule (Fig. 1C and fig. S1).

We next examined whether CPTX was equipped with the intended dual binding capacity *in vitro*. SPR assays showed that CPTX and NP1_PTX_ bound the ATD of GluA4 with comparable apparent affinities (*K*_D_ 4.5 ± 0.5 μM and 11.0 ± 0.9 μM, respectively; Fig. 1D and fig. S2). Like NP1_PTX_, CPTX also bound to the other GluA ATDs with a GluA3 > GluA1 > GluA2 preference. However, these interactions were weaker and accurate *K*_D_ values could not be reliably determined (Fig. 1D and fig. S2). In addition, CPTX specifically bound to the Nrx1β(+4) ectodomain (*K*_D_ 4.9 ± 0.9 nM), but not Nrx1β(–4) (Fig. 1E), thus retaining the strict isoform recognition specificity of Cbln1. Cell-based binding assays also showed that CPTX specifically bound human embryonic kidney (HEK) cells expressing GluA1–GluA4 ATDs, while Cbln1 only bound cells expressing the GluD2 ATD (fig. S3A). On the other hand, both Cbln1 and CPTX bound HEK cells expressing full-length Nrx1α, Nrx1β, Nrx2β, and Nrx3β only when the SS4 insert was present (fig. S3B). The ectodomain of Nrx1β(+4) specifically bound HEK cells expressing GluA4 ATD under CPTX application (fig. S3C). These results indicate that CPTX can simultaneously bind GluAs and neurexin isoforms containing the SS4 insert.

### CPTX induces excitatory pre- and postsynaptic sites *in vitro*

We examined whether CPTX could serve as a synaptic organizer with the designed GluA *vs* GluD specificity in neuronal cell cultures. Application of CPTX to *Cbln1*-null cerebellar granule cells induced accumulation of endogenous synaptophysin, a synaptic vesicle marker, on co-cultured HEK cells displaying GluA ATDs on their surface, but not on cells displaying the GluD2 ATD (Fig. 2A). Similarly, CPTX induced accumulation of synaptophysin in wild-type hippocampal neurons that contact HEK cells displaying GluA ATDs (fig. S4A). Conversely, application of Cbln1 induced presynaptic synaptophysin accumulation on co-cultured HEK cells expressing GluD2, but not GluA ATDs (Fig. 2A and fig. S4A). Beads coated with Cbln1 or CPTX accumulated presynaptic sites positive for endogenous synaptophysin, Nrx and vesicular glutamate transporter1 (VGluT1) (Fig. 2B and fig. S4C) in cultured hippocampal neurons, indicating that CPTX is capable of inducing presynapses *in vitro*. In contrast, NP1 bound HEK cells displaying the GluA4 ATD but failed to accumulate synaptophysin (fig. S4B).

**Fig. 2.**
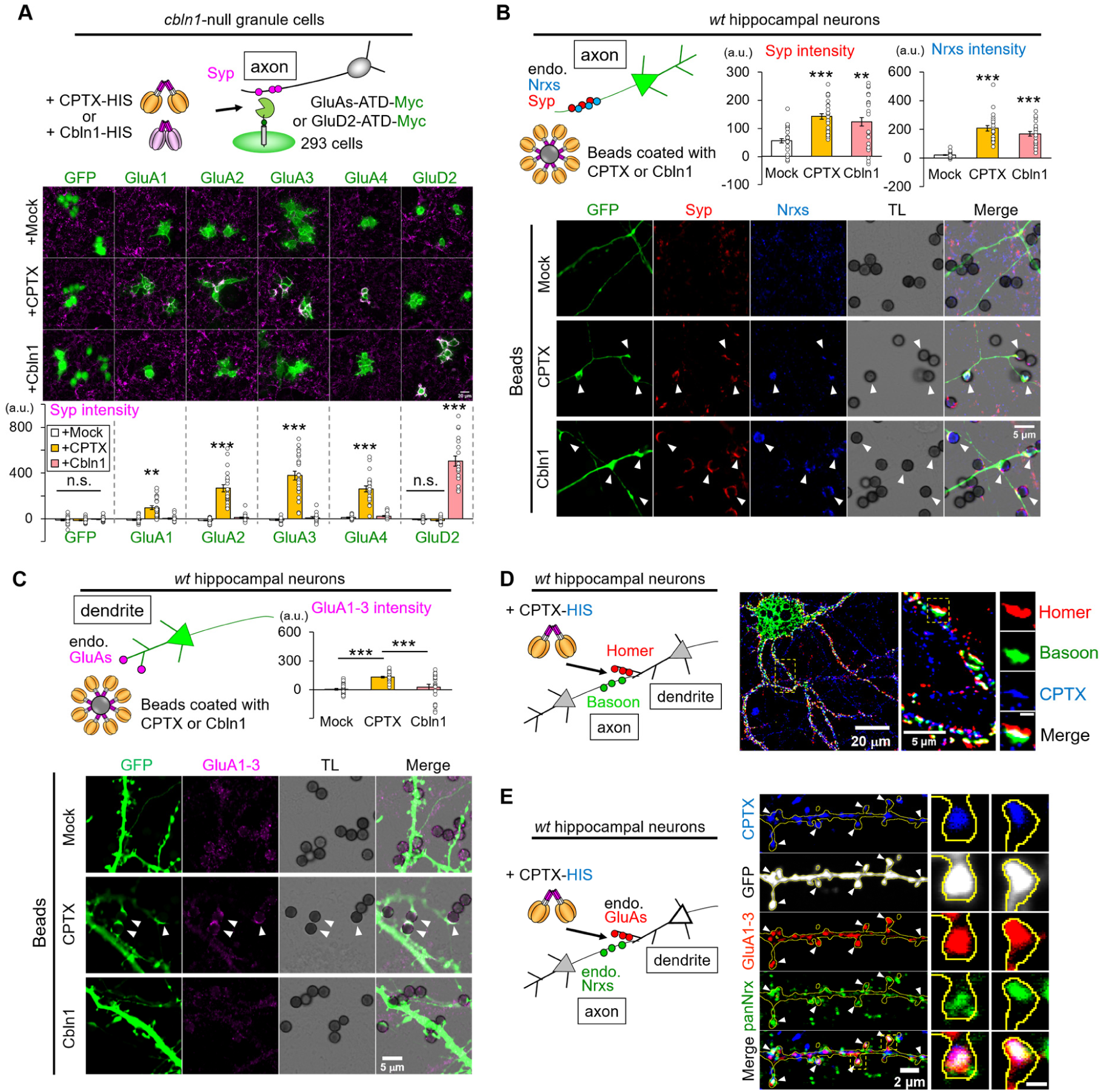
CPTX directly induces excitatory pre- and postsynaptic sites *in vitro*. (A) CPTX induces the accumulation of presynaptic terminals of cerebellar granule cells on co-cultured HEK cells displaying AMPA receptor ATDs. The intensities of synaptophysin immunoreactivity (Syp; magenta) were quantified on HEK cells (green) in the lower graph. Mock, vehicle (HEPES buffered saline (HBS)) controls. The bars represent mean ± SEM. \*\**P* < 0.01, \*\*\**P* < 0.001, *n* = 16–22 fields in 7 independent experiments, one-way ANOVA followed by Tukey’s test. Scale bar, 20 μm **(B)** Beads coated with CPTX or Cbln1 induce the formation of presynaptic boutons positive for endogenous synaptophysin (Syp) and neurexins (Nrxs) in co-cultured hippocampal neurons transfected with GFP. TL, transmitted light. Arrowheads indicate beads immuno-positive for Syp (red) and Nrxs (blue). Mock, beads coated with anti-HIS antibody but without CPTX or Cbln1. The intensities of Syp and Nrxs immunoreactivities were quantified on beads in the upper graphs. The bars represent mean ± SEM. \*\**P* < 0.01, \*\*\**P* < 0.001, *n* = 21 fields in 5 experiments, one-way ANOVA followed by Tukey’s test. Scale bar, 5 μm. **(C)** Beads coated with CPTX recruit endogenous GluA1–3 in dendrites of co-cultured hippocampal neurons. Arrowheads indicate beads immune-positive for GluA1–3 (magenta). Mock, beads coated with anti-HIS antibody as above. The intensity of GluA1–3 immunoreactivity was quantified on beads around dendrite in the upper graph. The bars represent mean ± SEM. \*\**P* < 0.01, \*\*\**P* < 0.001, *n* = 21 fields in 5 experiments, one-way ANOVA followed by Tukey’s test. Scale bar, 5 μm. **(D)** CPTX localizes at synapses. Representative immunocytochemical images show bassoon (green), homer (red) and HIS-tagged CPTX (blue) in hippocampal neurons. The boxed region is enlarged in the middle and the right panels. Scale bars, 20, 5 and 0.5 μm, respectively. (**E**) CPTX localizes between Nrxs and AMPARs. Representative images show immunoreactivity for pan-Nrxs (green), GluA1–3 (red), GFP (postsynaptic neuron, gray) and HIS-CPTX (blue) in hippocampal neurons. Arrows indicate dendritic spines immunopositive for Nrxs, GluA1–3 and CPTX. The boxed region is enlarged in the right panel. Scale bars, 2 and 0.5 μm, respectively.

We further asked whether CPTX could induce postsynaptic sites *in vitro*. Co-expression of Nrx1β(+4) and CPTX, but not Cbln1, on the surface of HEK cells facilitated accumulation of endogenous GluA1–GluA3 in dendrites of contacted hippocampal neurons (fig. S5A). Similarly, beads coated with CPTX, but not Cbln1, accumulated endogenous GluA1–GluA3 in dendrites of most hippocampal neurons (Fig. 2C) and GluA4 in parvalbumin-positive (PV^+^) interneurons (fig. S5B). Immunocytochemical analyses showed that soluble CPTX added to the culture medium accumulates between homer- and bassoon-positive puncta (Fig. 2D) and at the dendritic spines, accompanied by endogenous Nrx and GluA1–GluA3 (Fig. 2E and fig. S5C) in cultured hippocampal neurons. Together, these results indicate that CPTX serves as a bidirectional synaptic organizer by binding presynaptic Nrx(+4) and postsynaptic AMPARs *in vitro*.

### Rescue of synapse formation and motor coordination by CPTX in cerebellar ataxia mice

To test the impact of CPTX *in vivo*, we injected it into the cerebellum (lobules VI and VII) of adult *Cbln1*-null mice. Immunohistochemical analyses 1 d after the injection showed that CPTX was confined to the injected molecular layer (Fig. 3A) and colocalized with VGluT1 (a PF terminal marker), but not VGluT2 (a climbing fiber terminal marker) or the vesicular GABA transporter (a marker for inhibitory inputs) (Fig. 3B). This indicates that CPTX was mainly located at PF–PC synapses that express Nrx(+4) (*42*) and GluA1–GluA3 (*43*). As reported for Cbln1 (*13, 16*), single injections of CPTX were sufficient to partially restore morphological (fig. S6A) and functional (fig. S6B) PF–PC synapses and normal gait patterns (fig. S6C and fig. S7A; Movie S1 and Movie S2), as well as motor coordination measured by the rotor-rod test (fig. S7B) in the ICR mouse background, but not in the C57BL/6 one where the loss of skilled motor coordination phenotype was too severe. Similar to Cbln1 (*16*), the effect of CPTX on the rotor-rod performance recovery was maximal at 3 d post-injection but decayed afterwards (fig. S7C), suggesting that continued presence of CPTX is necessary to maintain PF–PC synapses in longer term. Together, these results indicate that CPTX could induce functionally normal PF–PC synapses in adult *Cbln1*-null mice.

**Fig. 3.**
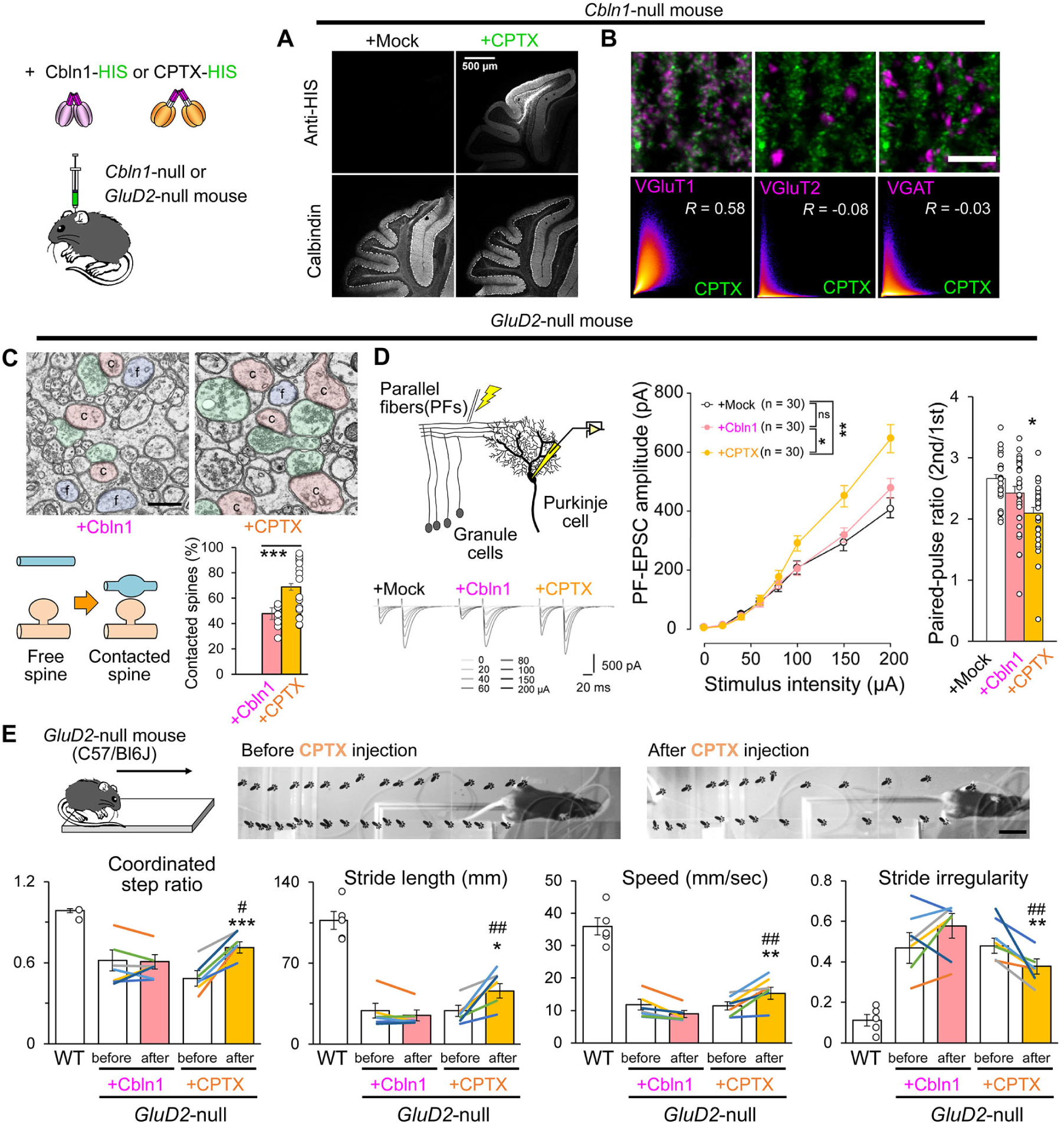
CPTX restores PF-PC synapses and motor coordination in *GluD2*-null mice. (**A, B**) HIS-tagged CPTX injected into *Cbln1*-null cerebellum localizes at PF–PC synapses. Immunohistochemical analyses showed HIS immunoreactivity in the molecular layer of the injected area (**A**) and colocalized with VGluT1 (a PF maker), but not VGluT2 (a climbing fiber marker) or VGAT (an inhibitory input marker). Mock (HBS) controls. (**B**). Scale bars, 500 μm (**A**) and 2 μm (**B**). The intensities of green and magenta signals in each pixel are plotted on x- and y-axis, respectively and Pearson’s correlation *R*-value is indicated in the lower panels. The degree of pixel density was shown as a heat map. (**C**) Representative electron microscopic images showing free dendritic spines (light blue) and contacted spines (light red) innervated by PFs (light green) in *GluD2*-null cerebellum 3 d after injection of Cbln1 or CPTX. Percentages of contacted PC spines are quantified in the lower graph. The error bars indicate the SEM. \*\*\**P* < 0.001, *n* = 10–20 sections from 1–2 mice, χ^2^ test. (**D**) CPTX restores functional PF-evoked EPSCs in *GluD2*-null mice 3 d after injection. Representative traces from slices prepared from *GluD2*-null mice that received HBS buffer (Mock), Cbln1 or CPTX are shown at the bottom left. The left graph shows averaged input-output relationship of PF–EPSCs for each treatment. n.s., not significant, \**P* < 0.05, \*\**P* < 0.01, *n* = 30 each, two-way ANOVA followed by Scheffe *post-hoc* test. The right graph shows the paired-pulse ratio of 2^nd^ to 1^st^ PF-EPSC amplitudes. \**P* < 0.05, \*\*\**P* < 0.001, *n* = 30 each, Kruskal-Wallis test followed by Scheffe *post-hoc* test. The error bars indicate the SEM. (**E**) CPTX improves the gait of *GluD2*-null mice 3 d after the injection. Representative footprints before and after CPTX injection are shown on the top. The lower graphs show the quantification of gait parameters. The bars are average scores and the lines are an individual score of each mouse before and after CPTX injection. \**P* < 0.05, \*\**P* < 0.01, \*\*\**P* < 0.001 compared with before values by Student’s paired *t*-test, ^#^*P* < 0.05, ^##^*P* < 0.01 compared with Cbln1 by Student’s *t*-test, *n* = 5 (WT), 6 (+Cbln1), 6 (+CPTX) mice.

We next investigated whether CPTX could serve as a synaptic organizer in the absence of GluD2, a Cbln1 receptor, *in vivo*. Electron microscopy analysis revealed that while Cbln1 was completely ineffective in *GluD2*-null mice 3 d after injection (*13*), CPTX partially restored PF–PC synapses (Fig. 3C). Whole-cell patch-clamp recordings in acute slice preparations also showed that, unlike Cbln1, CPTX increased the amplitudes of PF-evoked excitatory postsynaptic currents (PF-EPSCs) in *GluD2*-null mice (Fig. 3D) to the levels observed in wild-type mice (*13*). The increased paired-pulse ratio of PF-EPSC amplitudes, a phenotype indicative of decreased presynaptic release (*44*), was reset (Fig. 3D) to wild-type levels (*13*). Furthermore, unlike Cbln1, CPTX rescued the gait of *GluD2*-null mice (Fig. 3E and Movie S3). These results indicate that CPTX could induce morphologically and functionally normal PF–PC synapses independent of GluD2 *in vivo*.

### CPTX induces synaptic AMPAR accumulation in the wild-type hippocampus

To test if CPTX acts in brain regions outside the cerebellum, we injected CPTX into the dorsal hippocampus of wild-type mice. Super-resolution microscopy revealed that CPTX was localized between VGluT1 and AMPAR puncta in the CA1 *stratum radiatum* at 1 d after injection (Fig. 4, A to C and fig. S8B). Injected CPTX was also detected between bassoon-positive presynaptic terminals and GFP-positive postsynaptic sites of the CA1 pyramidal neurons in Thy1-GFP mice at 3 d after injection (fig. S8C). In contrast, CPTX did not show any colocalization with inhibitory synaptic markers (fig. S9). CPTX increased the number of strong GluA2/3-immunopositive signals, which was accompanied by PSD95 and VGluT1 signals (fig. S8A). Immunohistochemistry of CPTX and each AMPAR subunit (fig. S10A) revealed that CPTX shifted the intensity histogram of GluA1 to higher values, while causing re-distribution of GluA2/3 signals among puncta (fig. S10, B to D). As a result, CPTX increased the size of high-intensity (> mean + 3 SD) puncta of GluA1 and GluA2/3 (fig. S10, E and F). In contrast, while CPTX also shifted the intensity histogram of GluA4 to higher values (fig. S10D), it did not affect the size of high-intensity GluA4 puncta (fig. S10F). These results indicate that CPTX selectively binds to excitatory synapses in the hippocampus and modifies the distribution of AMPARs in a subunit-dependent manner.

**Fig. 4.**
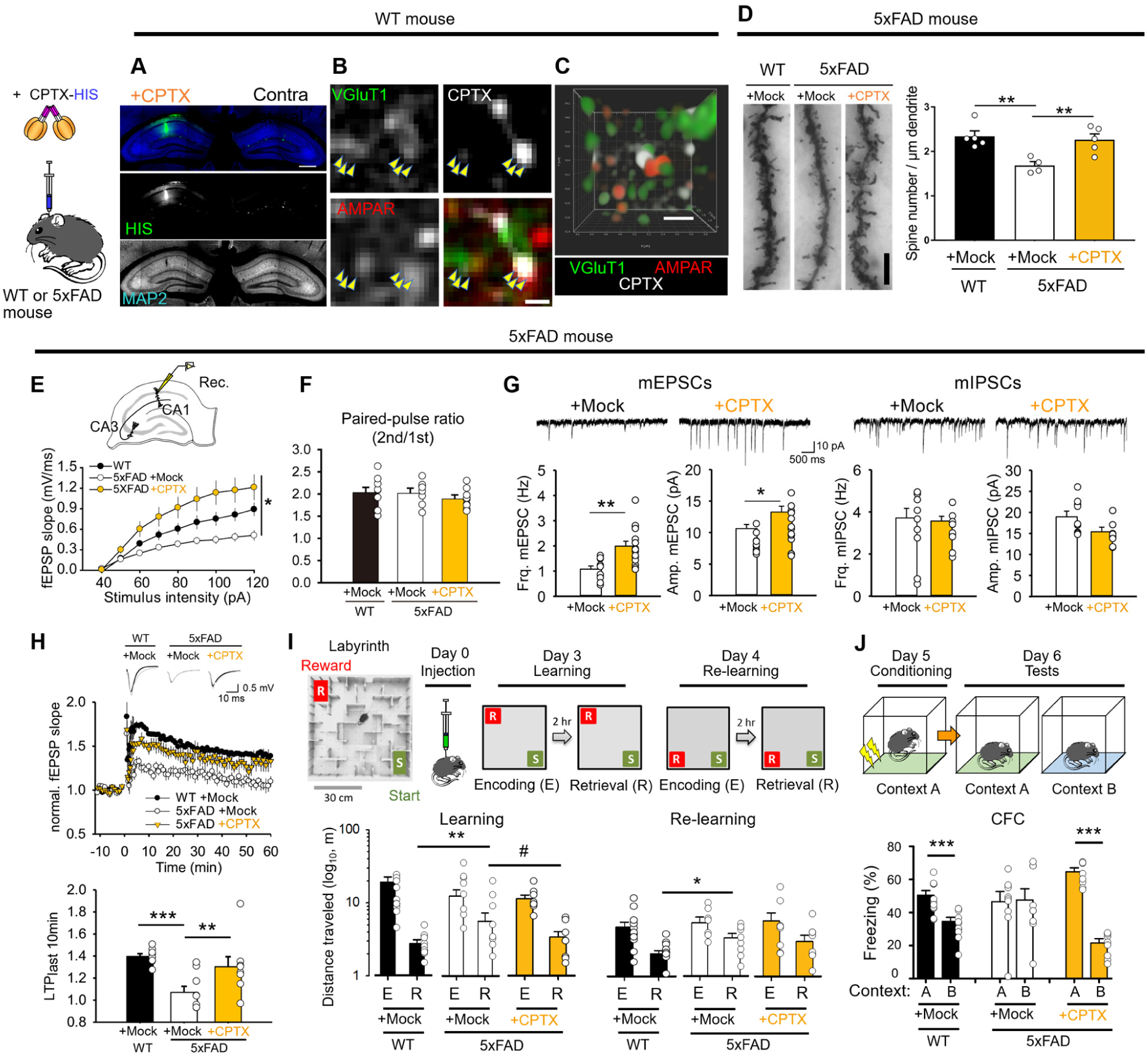
CPTX restores spines, LTP and hippocampus-dependent behaviors in Alzheimer’s disease model. (**A**-**C**) Injected CPTX localizes at excitatory synapses in the hippocampus of WT mice. (**A**) Representative immunohistochemical images of MAP2 (blue) and HIS-tagged CPTX (green) injected in the dorsal hippocampus at 1 d after the injection. The contralateral side is shown for comparison. Scale bar, 0.5 mm. (**B**) Airyscan super-resolution immunohistochemical images of the CA1 stratum radiatum of the hippocampus show CPTX (gray) in the proximity of VGluT1 (green) and AMPARs (red), marked by arrowheads. Scale bar, 0.2 μm. (**C**) 3D-reconstruction image showing CPTX localization between VGluT1- and AMPAR-positive sites. Scale bar for the x-axis, 0.5 μm. (**D**) CPTX restores spine density in 5xFAD mice. Representative Golgi-Cox staining of apical dendrites of CA1 pyramidal neurons in the dorsal hippocampus. Mock (HBS) controls. Scale bar, 5 μm. The graph shows averaged spine density of 7–8 secondary apical dendrites for each animal. The error bars indicate the SEM. \*\**P* < 0.01, *n* = 4–5 mice, one-way ANOVA followed by Holm-Sidak *post-hoc* test. (**E**) CPTX restored Schaffer collateral (SC)-evoked field excitatory postsynaptic potentials (fEPSPs) in 5xFAD mice. The graph shows averaged input-output relationship of SC–fEPSPs for each treatment. Mock, vehicle (HBS buffer) controls. The error bars indicate the SEM. \**P* < 0.05 compared with 5xFAD+Mock, *n* = 8 (WT), 9 (5xFAD+Mock), 8 (5xFAD+CPTX) slices, repeated two-way ANOVA. (**F**) CPTX does not affect the presynaptic release probability. The graph shows the paired-pulse ratio of 2^nd^ to 1^st^ SC-fEPSP amplitude at 50-ms intervals. The bars indicate the mean + SEM. \**P* < 0.05 compared with WT+Mock, *n* = 8 (WT+Mock), 9 (5xFAD+Mock), 8 (5xFAD+CPTX) slices, Student’s *t*-test. (**G**) CPTX increases the frequency and the amplitude of mEPSCs, but not mIPSCs, in 5xFAD hippocampus. \**P* < 0.05, \*\**P* < 0.01 compared with mock-treated control slices. *n* = 9–15, Student’s *t*-test. Representative traces are shown on the top. (**H**) CPTX restores LTP in 5xFAD mice. Application of theta-burst stimulation (TBS) to SC for three times induced a robust LTP at SC-CA1 synapses in WT, but not in 5xFAD hippocampus. Representative SC-fEPSP traces are shown on the top. The time course of LTP is shown by plotting the slope of SC-fEPSP against time. The error bars indicate the SEM. \*\**P* < 0.01, \*\*\**P* < 0.001, one-way ANOVA on ranks followed by Holm-Sidak *post-hoc* test. Also after exclusion of outlier in the CPTX-treated group, there are differences between all three groups (*P* < 0.05). (**I**) CPTX improves spatial memory in 5xFAD mice. CPTX or mock, vehicle (HBS buffer) was injected in 5xFAD mice on day 0. On day 3, mice were placed at the start point (S) of a 3D-printed maze and the pellet was placed at the reward point (R). Two hours after the initial encoding (E) session, mice were returned to the same start point to examine memory integrity in the retrieval (R) session. On the next day, the position of reward was changed, and reversal learning was evaluated. The averaged total distances that mice traveled to reach the goal during the encoding and the retrieval sessions are shown in the lower graph. Log_10_ scaling of Y-axis is used to facilitate comparison of distances before and after training. The error bars indicate the SEM. ^#^*P* < 0.1, \**P* < 0.05, \*\**P* < 0.01, two-way RM ANOVA followed by LSD Fisher’s *post-hoc* test. *n* = 8–11 mice. (**J**) CPTX restores contextual fear conditioning performance in 5xFAD mice. CPTX or vehicle (Mock) was injected in 5xFAD mice on day 1. Electrical shock was applied to wild-type and 5xFAD mice in context A on day 5. Freezing time was measured in context A and context B on day 6. The lower graph shows mean (± SEM) freezing time of each group. \*\*\**P* < 0.001, Student’s paired *t*-test.

Functionally, the CPTX injection increased the frequency of miniature EPSCs (mEPSCs) in CA1 pyramidal neurons as early as 4 h after application to acute hippocampal slices (fig. S11A). CPTX also upregulated both the amplitude and frequency of mEPSCs in hippocampal slices prepared 3 d after injection (fig. S11B), but did not enhance the frequency and amplitude of mEPSCs in CA1 PV^+^ interneurons (fig. S12), which express high levels of GluA4 and NP1/NPR (fig. S13A and B). CPTX injections showed stronger effects in increasing the GluA1–3 intensity in the *lacunosum moleculare* (fig. S14B vs. fig. S10B) where NP1/NPR immunoreactivities were much weaker than in the *stratum radiatum* (fig. S13G). These results indicate that CPTX can rapidly induce functional excitatory synapses, but its impact depends on the endogenous levels of NPs and AMPAR subtypes, presumably due to the competition with endogenous proteins. For example, the ATDs of GluA4, where CPTX binds, would most likely be occupied by endogenous NPs in PV^+^ interneurons.

### CPTX restores spines, LTP and cognition in an Alzheimer’s disease model

Prompted by these observations, we next examined whether CPTX can modify neuronal circuits in the hippocampus under pathological conditions *in vivo*. We used 5xFAD mice, a model of early familial Alzheimer’s disease (AD) which allowed us to test the impact of CPTX in the context of severe loss of neurons, synapses, synaptic plasticity and cognitive abilities (*45, 46*). Golgi staining revealed reduced spine densities in the CA1 region of middle-aged 5xFAD mice (15–18 months old; Fig. 4D). Extracellular field recordings in the CA1 region confirmed that excitatory postsynaptic potentials (fEPSPs) induced by Schaffer collateral (SC) stimulation were reduced in acute slices prepared from 5xFAD mice (Fig. 4E) as reported earlier (*47*). Injection of CPTX into the hippocampus region of 5xFAD mice restored spine density (Fig. 4D) as well as the amplitude of SC-fEPSPs, which was even larger than in the wild-type hippocampus (Fig. 4E). In contrast, CPTX did not change the paired-pulse facilitation ratio of fEPSPs (Fig. 4F). Moreover, whole-cell patch-clamp recordings from CA1 pyramidal neurons revealed that the frequency and the amplitude of mEPSCs were increased in 5xFAD mice injected with CPTX (Fig. 4G). On the other hand, the frequency and the amplitude of miniature inhibitory synaptic currents (mIPSCs) were unaffected (Fig. 4G). These results indicate that CPTX specifically improved excitatory transmission between principal cells without changing the presynaptic properties or GABAergic transmission in 5xFAD mice.

Impaired long-term potentiation at SC-CA1 synapses (SC-LTP) is believed to underlie certain cognitive abnormalities in 5xFAD mice (*47*). Interestingly, CPTX injection rescued impaired SC-LTP in middle-aged 5xFAD mice (Fig. 4H), which may reflect restored spines (Fig. 4D) capable of synaptic plasticity. To evaluate spatial memory in a paradigm with minimal stress exposure, we placed three groups of mice in a labyrinth with food pellets as a reward and measured the total distance that mice traveled to reach the goal during the encoding and retrieval sessions (Fig. 4I). Although there was no difference in the travel distance between each group during the encoding session, 5xFAD mice traveled longer distance than wild-type mice during the retrieval session. CPTX injection 3 d before the learning test decreased the distance that 5xFAD mice traveled in the retrieval session, suggesting an improvement of spatial learning after the CPTX treatment. After reversal learning, when mice had to find a reward in a new position, the performance of 5xFAD mice was impaired in the mock, but not in CPTX-injected group, as compared to wild-type mice (Fig. 4I). Next, we evaluated conditioned fear memory by exposing the same three groups of mice (wild-type mice injected with mock (buffer), 5xFAD mice injected with mock or CPTX) to an electrical shock in context A and measuring the freezing time in the conditioned context A and a neutral context B 1 d later. 5xFAD mice failed to discriminate context A from B, i.e. showed generalized rather than context-specific learned fear response. CPTX injection 5 d before conditioning greatly improved the performance of 5xFAD mice in the contextual fear conditioning paradigm, so they showed context discrimination even better than wild-type controls (Fig. 4J). Together, these results indicate that the impaired SC-LTP and hippocampus-dependent learning in middle-aged 5xFAD mice were rescued by the injection of CPTX.

### CPTX restores synapses and locomotion in spinal cord injury models

Re-organization and the E/I balance of spared intraspinal networks have recently been shown to contribute to functional recovery after spinal cord injury (SCI) (*48, 49*). Thus, to test whether CPTX could restore glutamatergic circuits in the spinal cord, we performed dorsal hemisections at the thoracic level 10 (T10) in mice and injected CPTX rostral to the injured region (Fig. 5A). Spinal cords were dissected 2–5 d after injections and coronal serial sections were immunostained for VGluT2 and CPTX (fig. S15A). CPTX was distributed in the gray matter and was largely colocalized with VGluT2 and GluA4 immunopositive (VGluT2+ and GluA4+) puncta in a region 1.4–1.6 mm rostral to the epicenter of the injured site (Fig. 5B and fig. S15B). Super-resolution microscopy revealed that CPTX was located between VGluT2+ and GluA4+ puncta (Fig. 5C). Since most excitatory spinal cord interneurons that are involved in locomotion express high levels of VGluT2 and GluA4 (*50*), these results indicate that CPTX may connect excitatory pre- and postsynaptic sites of spared neuronal circuits.

**Fig. 5.**
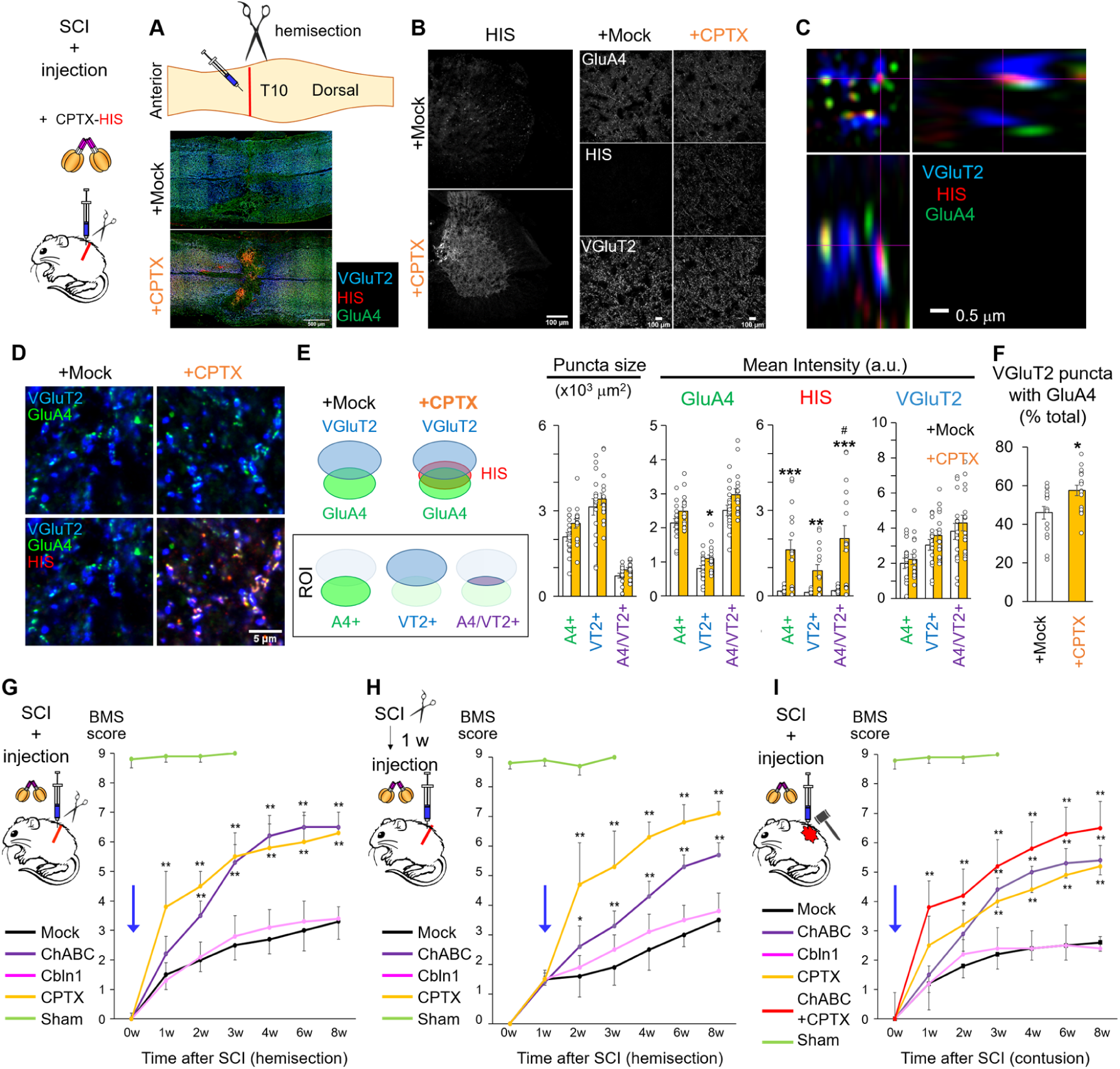
CPTX promotes restoration of synapses and motor function in spinal cord injury. (**A**) Schematic depiction of SCI caused by hemisection at T10. Lower panels indicate representative horizontal sections of spinal cord immunostained for VGluT2, HIS (CPTX) and GluA4 upon mock (HBS) or CPTX injection. (**B**) Representative coronal sections at approximately 1.5 mm from the injury epicenter immunostained for VGluT2, HIS and GluA4 upon mock (HBS) or CPTX injection. Right panels show high-magnification views in the dorsal region. Scale bars, 100 μm. (**C**) A representative orthogonal image obtained by Airyscan super-resolution microscopy indicating the localization of CPTX in the proximity of VGluT2- and GluA4-immunopositive puncta. Scale bar, 0.5 μm. (**D–F**) Quantification of excitatory synapses induced by CPTX. Representative immunohistochemical images of coronal sections stained for VGluT2 (blue), GluA4 (green) and HIS (red; CPTX) in the HBS buffer (Mock) or CPTX injection (**D**). Scale bar, 5 μm. The size and the signal intensity of GluA4, VGluT2 and HIS were quantified in each region of interest (ROI; GluA4+ (A4+), VGluT2+ (VT2+) or GluA4+/VGluT2+ (A4/VT2+)) (**E**). The percentage of VGluT2+ puncta that were accompanied with GluA4+ puncta is also shown (**F**). \**P* < 0.05, \*\**P* < 0.01, \*\*\**P* < 0.001 compared with mock by Student’s *t*-test, ^#^*P* < 0.05 compared with GluA4 or VGluT2 by Student’s paired *t*-test, *n* = 16 slices each for mock and CPTX from 8 mice. (**G–I**) Time-course analyses of locomotion (BMS score) in SCI mice after the injection. Mock (HBS buffer vehicle control), ChABC, Cbln1 or CTPX was injected into the spinal cord immediately (**G**) or 1 week (**H**) after hemisection, or immediately after contusion by impactors (**I**, 90 kdyn). For sham controls, the spinal cord was surgically exposed without imposing injury or injection. Mice that showed BMS score = 1.5 at 1 week after hemisection were selected for H. \**P* < 0.05, \*\**P* < 0.01, \*\*\**P* < 0.001 compared with mock and ^#^*P* < 0.05, \*\**P* < 0.01 compared with ChABC by repeated two-way ANOVA with *post-hoc* Bonferroni-Dunn test, *n* = 9 mice for each treatment.

We next assessed whether CPTX induced excitatory synapses by quantifying the intensity and the size of VGluT2+, GluA4+ and CPTX+ puncta (Fig. 5D). CPTX increased the size of GluA4+ and GluA4+/VGluT2+ (double positive) puncta, as well as GluA4 signal intensities in VGluT2+ puncta (Fig. 5E). Furthermore, CPTX increased the percentage of VGluT2+ puncta that were also GluA4+ (Fig. 5F). These results indicate that CPTX induced GluA4+ excitatory synapses within regions spared by the injury (fig. S15A and fig. S15B).

Finally, we assessed the recovery of locomotion after spinal cord hemisections. For comparison, we injected separately chondroitinase ABC (ChABC), an enzyme that promotes robust nerve regeneration (*51*), and Cbln1 to the lesioned spinal cord. During the acute stage, single injections of CPTX as well as ChABC, but not Cbln1, significantly restored the Basso Mouse Scale (BMS) score (*52*) (Fig. 5G and Movie S4) and other locomotive parameters examined until 6 weeks after the injury (fig. S15D and Movie S5). ChABC delivered 1 week after the hemisection was less effective (Fig. 5H), as reported previously (*53*). In contrast, when CPTX was delivered 1 week after injury it was most effective (Fig. 5H, fig. S15C and Movie S6 and 7). CPTX also efficiently restored locomotion in a contusion SCI model, which is considered to better mimic the pathophysiology of naturally occurring injuries in people (Fig. 5I and fig. S15E). In this paradigm, a combination of CPTX plus ChABC treatment showed the most prominent recovery of locomotion, especially in the early stages after contusion. Together, these results indicate that CPTX promotes functional recovery in spinal cord injury models by enhancing excitatory connectivity in spared neuronal circuits. Mechanistically, this process appears to be distinct from the one induced by ChABC treatment.

### Discussion

We demonstrate that a synthetic, structure-guided, synaptic organizer termed CPTX can induce functional and structural synapses in the cerebellar, hippocampal and spinal cord neuronal circuits *in vivo*. CPTX increased the frequency of mEPSCs as early as 4 h, and then both the frequency and the amplitude of mEPSCs 3 d after application to the wild-type hippocampus (fig. S11) without affecting presynaptic release probability (Fig. 4F). Thus, we propose that CPTX induces functional synapses by connecting pre-synaptic neurexin with post-synaptic AMPARs and facilitating trans-synaptic alignment of AMPARs with the synaptic vesicle release sites. Such “molecular bridges” may initiate and integrate within the trans-synaptic nanocolumns described at excitatory synapses (*40*). Interestingly, CPTX accumulated AMPARs by increasing GluA1 intensity in all puncta and causing re-distribution of GluA2/3 signals in the hippocampus 1 d after injection (fig. S10). Insertion of new GluA1 subunits to plasma membranes is thought to be a crucial step for spine enlargement during LTP (*54*); therefore, we propose that CPTX induces structural changes by clustering GluA1, and possibly other associated proteins, at Nrx(+4)-positive sites. Since molecular components are considerably different among neuronal synapse types and circuits, further studies are warranted to test this hypothesis and clarify the detailed molecular mechanisms by which CPTX restores functional and structural excitatory synapses under pathological conditions.

Novel ESPs continue to be discovered. For example, LGI1, an ESP produced by the hippocampal neurons, is particularly interesting because it indirectly recruits AMPARs by binding to pre- and postsynaptic ADAM23 and ADAM22, respectively (*55, 56*). Since ESPs generally have a modular structure, we anticipate that a toolkit of synthetic ESPs with a variety of pre- and postsynaptic specificities, able to restore or modify synaptic connectivity in various neuronal circuits, could be developed. The structure-guided design of ESPs that target specific GABA_A_ receptor heteromers (*57*), as well as specific AMPAR or KAR subunits, will be useful to restore E/I balance in specific neuronal circuits affected by autism spectrum disorders and schizophrenia (*1, 5, 58*), epilepsy (*3*) and Alzheimer’s disease (*2*). Importantly, while the effect of CPTX was transient in *Cbln1*-null ataxia mice (fig. S7C), a single injection of CPTX greatly enhanced locomotion for at least 6-7 weeks in SCI models (Fig. 5, G and H), suggesting that the effect of synthetic ESPs could last very long under certain conditions, probably by recruiting additional, endogenous, synaptic organizers. Thus, synthetic ESPs built upon the principles described here can become powerful tools to investigate and potentially cure disorders associated with impaired neuronal connectivity.

## Acknowledgments

We thank staff at the Diamond Light Source beamline I03, T. Walter and K. Harlos for support with protein crystallization, E. Budinger, O. Kobler and J. Schneeberg for histology advice and assistance. This work was funded by the Human Frontier Science Program (grant RGP0065/2014 to M.Y., A.D. and A.R.A.), the UK Medical Research Council (MRC) (grants L009609 and MC_UP_1201/15 to A.R.A.), the Japan Society for the Promotion of Science (JSPS) (grant 15H05772 and 16H06461 to M.Y., 17K10949 and 1705584 to K.T., 25893233, 26860148, 14J07587 and 18K19380 to K.S., 18H04563 to W.K.), the Keio University Grant-in-Aid for Encouragement of Young Medical Scientists (to K.S.), the Keio Association Grant-in-Aid (to K.S.), the Astellas Foundation for Research on Metabolic Disorders (to K.S.), the Daiichi Sankyo Foundation of Life Science (to K.S.), the AMED (iD3 16nk0101302h0002 to K.T. and JP18dm0107124 to W.K.), the JST CREST (JPMJCR1854 to M.Y.), the JST ERATO (JPMJER1802 to W.K.), the Takeda Science Foundation (to M.Y. and W.K.), the Marie-Curie Actions postdoctoral fellowship (grant 328531 to J.E.), the University of Bordeaux Initiative of Excellence (IdEx) fellowship (to J.E.), the European Research Council (ERC) (Starting Grant 850820 to J.E.), the European Union 7^th^ Framework Programme Initial Training Network (EU FP7 ITN) (grant 606950 EXTRABRAIN to A.D.), and the Bundesministerium für Bildung und Forschung (BMBF) (EnergI project, TP5, 01GQ1421A to A.D.).

## Author Contributions

K.S. performed cell biological experiments, immunohistochemical experiments of hippocampus, cerebellum and spinal cord tissues, behavioral studies of cerebellar function, analyzed the data and wrote the paper. A.J.C., J.E. and A.R.A. collected X-ray data and solved the NP1_PTX_ structure; A.R.A. conceived the CPTX organizer; J.E. performed SPR and MALS experiments; W.K. carried out electrophysiological analyses in the cerebellum; I.S. performed electrophysiological experiments in the hippocampus; J.E., V.T.C. and A.J.C. prepared recombinant proteins; O.S. developed 3D printed labyrinth and performed behavioral studies of hippocampal function; M.F.F. and S.S. performed spine analysis; E.M. performed electron microscopic analyses; R.K. performed immunocytochemical and immunohistochemical analyses of hippocampal neurons; E.M. performed electron microscopic analysis; K.S. and T.S. performed cell-based binding assays; S.O. developed analytical tools for gait performance; H.S., M. I., Y.M., Y.T. and K.T. generated the experimental models of spinal cord injury and performed behavioral analyses. M.W. raised new antibodies against neuronal pentraxins. A.D., A.R.A. and M.Y. designed and supervised the project and wrote the paper. All authors provided feedback on the final manuscript version.

## Materials and Methods

### Mice

Wild-type (Charles River, US and Japan SLC, Inc) and *Cbln1*-null mice were used for primary culture. In experiments assessing cerebellar functions, we used male and female *Cbln1*-null mice (*1*) with C57Bl/6J or ICR background and *GluD2*-null mice (*2*) with C57Bl/6J. They were housed on a conventional 12 h light/dark cycle (light on at 8:30 a.m.). The behavioral tests were conducted in the light phase of the cycle. In experiments assessing hippocampal functions, we used male C57Bl/6J, PV-*Cre*/tdTomato double transgenic mice (Pvalb ^tm1(cre)Arbr^, 017320, Jackson Laboratory/tdTomato Ai9) (*3*), and 5xFAD mice, which carry 5 familial Alzheimer’s disease mutations (*4*), and their wild-type littermates with C57Bl/6J background (Charles River, US). At least one week before starting the experiments, mice were transferred from the main animal facility of DZNE (Magdeburg, Germany) to a small vivarium, where they were housed individually with food and water *ad libitum* before food restriction diet (FRD) was given, on a reversed 12 h light/dark cycle (light on at 9:00 p.m.). All behavioral experiments assessing hippocampal functions were performed in the late afternoons during the dark phase of the cycle when mice were active, under constant temperature (22 ± 1°C) and humidity (55 ± 5%). For the spinal cord injury (SCI) model, wild-type mice with ICR background (Japan SLC, Inc) were used. They were housed on a conventional 12 h light/dark cycle. Locomotor functions were assessed at a fixed time in the evening. All treatments and behavioral procedures were conducted in accordance with ethical animal research standards defined by German and Japanese law and approved by the Ethical Committee on Animal Health and Care of the State of Saxony-Anhalt, Germany, with license numbers 42502-2-1343 DZNE and 42502-2-1322 DZNE, by the Animal Resource Committee of Keio University (No. 09050) and by the Institutional Animal Care and Use Committee of Aichi Medical University (No. 1559).

### Plasmids

For surface plasmon resonance (SPR) experiments, cDNAs encoding the extracellular human glutamate receptor D2 amino-terminal domain (ATD; UniProt ID O43424; GluD2 ATD: Asp24-Gly440), the mouse glutamate receptor A1 ATD (UniProt ID P23818; GluA1 ATD: Ala19-Thr412), the mouse glutamate receptor A2 ATD (UniProt ID P23819; GluA2 ATD: Val22-Thy419), the mouse glutamate receptor A3 ATD (UniProt ID Q9Z2W9; GluA3 ATD: Gly23-Thr422), the mouse glutamate receptor A4 ATD (UniProt ID Q9Z2W8; GluA4 ATD: Gly21-Thr420), the human β-neurexin-1 LNS6 domain (GenBank ID NM_138735; β-Nrx1 LNS6: His85-Val265) and the pentraxin domain of human neuronal pentraxin-1 (NP1_PTX_; UniProt ID Q15818; Pro224-Ile431) were fused to the C-terminus of a hexa-histidine (His6) tag and cloned into the pHLsec expression vector (*5*). To produce a trimeric form of the pentraxin domain, NP1_PTX_ was fused to the C-terminus of the three-stranded GCN4 leucine zipper coiled-coil sequence (*6*), to yield NP1_PTX-3COIL_ (Fig. 1C). For large-scale protein production, constructs cloned in the pHLsec vector were used for the transient expression. For CPTX production, a lentivirus expression system was also employed, by subcloning the cDNA into the pHR-CMV-TetO_2_ vector and co-transfecting it with the psPAX2 packaging plasmid and the pMD2.G envelope plasmid (*7*). To produce CPTX, NP1_PTX-3COIL_ was fused to the C-terminus of the cysteine-rich region (CRR) (*8*) of human Cbln1 (UniProt ID P23435; Gln22-Ile53) (Fig. 1C). For the *in vitro* binding and synapse formation assays, mouse NP1-full length (GenBank ID NM_008730.2) was cloned into pCAGGS vector (kindly provided by Dr. J. Miyazaki, Osaka University, Osaka, Japan) with a C-terminal HA-tag, β-neurexin-1 full-length cDNA was cloned into pCAGGS with a C-terminal FLAG-tag, β-neurexin-1 extracellular domain was cloned into the pCAGGS vector with a C-terminal human Fc tag and the ATD regions of GluA1–4 were cloned into a modified pDisplay vector (Invitrogen), as described previously (*9*). The miR sequences were inserted upstream of IRES-GFP (green fluorescent protein) in the pCAGGS vector, as described previously (*10*). The 21-bp target sequences of mouse *Nptx1* and *Nptxr* genes were designed by BLOCK-iT (Invitrogen) as follows: AGA CAA GTT TCA GCT GAC ATT (for NP1) and TGC TCA GTC GCT TCC TCT GTA (for NPR). For immunocytochemical analyses to check the selectivity of anti-NPs antibodies, mouse NP1-full length (GenBank ID NM_008730.2), NP2-full length (GenBank ID NM_016789.3) and NPR-full length (GenBank ID NM_030689.4) were cloned into the pCAGGS vector with a C-terminal HA-tag. For the immunoblotting analyses to check the selectivity of anti-NPs antibodies, mouse NP1 (without signal sequence, aa 1-22), NP2 (without signal sequence, aa 1-14) and NPR (without transmembrane domain, aa 1-23) were cloned into pCAGGS downstream of the Igκ secretion signal sequence, followed by a N-terminal 2xHA tag.

### Preparation of Recombinant Proteins

For large-scale protein production, proteins were expressed transiently (*5*) or lentivirally (*7, 11*). For transient transfections, the expression vectors were transfected into HEK293T (ATCC no. CRL-1573) (in the presence of the class I α-mannosidase inhibitor kifunensine (*12*)) or HEK293S-GnTI^-/-^ (*13*) cells, using an automated procedure (*14*). Five (HEK293T) to ten (HEK293S-GnTI^-/-^) days post-transfection, the conditioned medium was collected and filtered (0.22 μm) for protein purification. For the lentiviral infection, the proviral DNA, packaging and envelope plasmid were co-transfected to HEK293T using PEI. The supernatant containing viral particles was harvested after 2–3 days, passed through 0.45 mm filters and directly used for the infection to HEK293T cells. The infected cells were grown and expanded to 3 L cultures. Supernatants containing secreted recombinant proteins were collected 7-10 days post-infection and filtered (0.22 μm) prior to protein purification. The collected supernatants were concentrated and buffer-exchanged using the AKTA-flux system (GE Healthcare) or a QuixStand benchtop diafiltration system. Proteins were purified by immobilized metal-affinity chromatography (IMAC) using pre-packed nickel Sepharose columns (GE Healthcare), or in batch mode on TALON^®^ beads (Takara Bio) collected on an EconoColumn (Bio-Rad). Eluting fractions containing the proteins of interest, according to SDS-PAGE analysis, were pooled and concentrated, followed by the further purification by size exclusion chromatography using a Superdex 200 16/600 column (GE Healthcare) in 10 mM HEPES (4-(2-hydroxyethyl)-1-piperazineethanesulfonic acid) pH 7.4, 150 mM sodium chloride and 3 mM calcium chloride (HBS-C) for structural studies, or in 10 mM HEPES, pH 7.4, 150 mM sodium chloride (HBS) for biological studies, or in 10 mM Tris pH 7.4, 150 mM sodium chloride, 3 mM calcium chloride and 0.005% (v/v) Tween-20 (TBS-CT) for interaction studies using SPR.

### Gene Splicing and Site-Directed Mutagenesis

A multiple-step overlap-extension PCR (Pyrobest Polymerase, Takara Bio) was used for site-directed mutagenesis and construction of chimeric protein constructs; the resulting PCR products were cloned into the pHLsec-His6, pHLsec-Avitag3, or derived vectors (*5*). The following internal primer pair was used for the introduction of human Nrx1β spliced sequence #4 (SS4; GNNDNERLAIARQRIPYRLGRVVDEWLLDK) into Nrx1β(–4): 5’-CGC ATT CCC TAT CGG CTA GGG AGA GTG GTG GAC GAA TGG CTG CTC GAT AAA GGG AGG CAA CTG ACC ATC TTC AAC TCA C-3’; 5’-CCC TAG CCG ATA GGG AAT GCG TTG CCG TGC TAT GGC TAA CCT CTC ATT GTC GTT GTT TCC AGC TGG GTA TCT CTC AAT GAC-3’

### Protein Crystallization

Crystallization trials, using 100 nL protein solution plus 100 nL reservoir solution in sitting drop vapor diffusion format, were set up in 96-well Greiner plates using a Cartesian Technologies robot (*15*). Purified human NP1_PTX_ (Pro224-Ile431), concentrated to 6.5 g/L, crystallised in 15% (v/v) glycerol, 25.5% (w/v) polyethylene glycol (PEG) 8000, 0.17M ammonium sulphate and 0.085 M sodium cacodylate pH 6.5. Crystals were cryoprotected using a reservoir solution containing 30% (v/v) ethylene glycol.

### Crystallographic Data Collection and Structure Determination

Diffraction data for NP1_PTX_ were collected at Diamond Light Source (DLS) beamline I03 to a nominal resolution of 1.45 Å in space group (SG) C2. All data were indexed, integrated, and scaled using the automated XIA2 expert system (*16*), using the Labelit (*17*), POINTLESS and AIMLESS (*18, 19*), and XDS (*20*) programs. The structure of NP1_PTX_ was solved by molecular replacement using the program Phaser (*21*) and using the human SAP (Serum Amyloid P Component; PDB code 1SAC) crystal structure (*22*) as a search model. Crystallographic data collection and refinement statistics are presented in Table S1.

### Crystallographic Refinement and Model Analysis

Maximum-likelihood refinement of NP1_PTX_ was initially performed with Refmac using “jelly body” restraints (*23*), and finally with the PHENIX suite (*24*), with automated X-ray and atomic displacement parameter (ADP) weight optimization applied throughout. Automated model building was performed using ARP/wARP (*25*) and further manual model building was performed using Coot (*26*). Structure validation was performed with the PHENIX program suite using MolProbity routines (*24, 27*). Molecular representations were made using the program PyMol (*28*).

### Multi-angle Light Scattering (MALS)

Protein samples concentrated to ∼1.0 g/L were injected into an HPLC-driven SEC column (Superdex 200 10/30 column, GE Healthcare) equilibrated with HBS-C buffer. The SEC column was coupled to an online UV detector (Shimadzu), an 18-angle light scattering detector (DAWN HELEOS), and a refractive index detector (Optilab T-rEX) (Wyatt Technology). Proteins for MALS contained N-linked oligomannose-type sugars, and molecular mass determination was performed using an adapted RI increment value (dn/dc standard value; 0.185 mL/g) to account for the glycosylation state. Data analysis was carried out using the ASTRA V software (Wyatt Technology).

### Surface Plasmon Resonance (SPR)

cDNA for the immobilized proteins was cloned into the pHLsec-Avitag3 vector (*5*), resulting in proteins carrying a C-terminal biotin ligase (BirA) recognition sequence (Avitag). Constructs were co-transfected with pDisplay-BirA-ER (Addgene plasmid 20856; coding for an ER-resident biotin ligase) (*29*) for in vivo biotinylation in HEK 293T cells in small-scale 6- or 12-well plates in a 3:1 pHLsec:pDisplay stoichiometric ratio. A final concentration of 100 µM D-biotin was maintained in the expression medium to ensure near-complete biotinylation of the recognition sequence. After 48 h of expression, conditioned medium was collected and dialyzed against 10 mM Tris pH 7.4, 150 mM sodium chloride, 3 mM calcium chloride and 0.005% (v/v) Tween-20 (TBS-CT). SPR experiments were performed on a Biacore T200 machine (GE Healthcare) operated at a data collection frequency of 10 Hz; i.e. a temporal resolution of 0.1 sec. Streptavidin (Sigma-Aldrich) was chemically coupled via amine coupling chemistry onto CM5 chips to a response unit (RU) level of 5000 RU. Then, biotinylated proteins were captured to the desired RU level. In each instance, for every two analyte binding cycles, a buffer injection was performed, allowing for double referencing of the binding responses (*30*).

### Antibodies

The origin, dilution, company and catalog number are described as follows: Anti-calbindin (Goat, 1:500, Frontier Institute, Af1040), anti-HIS (Mouse, 1:1000, MBL, D291-3 or Rabbit, 1:1000, CST, #2365), anti-Myc (Rabbit, 1:1000, MBL, 562), anti-FLAG (Rabbit, 1:1000, Sigma-Aldrich, F7425), anti-HA (Mouse, 1:1000, BAbCO, MMS-101P), anti-synaptophysin (Mouse, 1:500, Sigma-Aldrich, S5768 or G, 1:500, Frontier Institute, Af300), anti-neurexin (Chicken, 1:500, a gift from Peter Scheiffele (*31*), anti-GFP (Rabbit 1:1000, Frontier Institute, Af-2020Frontier Institute), anti-GluA1 (Rabbit, 1:100, Calbiochem, PC246 or Guinea Pig, 1:500, Frontier Institute, Af380), anti-GluA2/3 (Rabbit, 1:1000, Chemicon, AB1506), anti-GluA4 (Rabbit, 1:1000, Pharmingen, 60666N or Guinea Pig, 1:500, Frontier Institute, Af640), anti-VGluT1 (Rabbit, 1:500, Frontier Institute, Af570 or Goat, 1:500, Frontier Institute, Af310), anti-VGluT2 (Guinea Pig, 1:500, Frontier Institute, Af720 or Goat, 1:500, Frontier Institute, Af310), anti-VGAT (Goat, 1:500, Frontier Institute, Af620)), anti-Parvalbumin (Goat, 1:500, Frontier Institute, Af460), anti-Homer 1 (Guinea pig, 1:1000, Synaptic Systems, 160-004) and anti-Bassoon (Rabbit, 1:500, Synaptic Systems, 141-003). Anti-NP1 and NPR were newly raised by guinea pig immunized with the peptide (NP1: aa 185-227, NPR: aa 226-263). For the simultaneous detection of various GluAs (i.e. GluA1–3 or GluA1–4), the mixture of the primary antibodies described above was used. Secondary antibodies conjugated with DyLight 405, Alexa 488, 546, 647 and Cy3 (Invitrogen or Jackson Lab) against the respective primary antibody were used in the dilution (1:1000 for immunocytochemistry, 1:200 for immunohistochemistry).

### Immunoblotting

Cells and conditioned medium were solubilized by laminae buffer (2% SDS, 80 mM Tris-HCl pH 6.8, 10% glycerol, 0.00625% Coomassie blue G 250) and the proteins were reduced by boiling for 3 min under 2% 2-mercaptoethanol. Samples were subjected to SDS-PAGE of gradient gel (Wako) and blotted onto the membrane (Millipore). Membranes were blocked with TS-tween (0.1% Tween20, 50 mM Tris-HCl pH7.6, 150 mM NaCl) containing 5% skim milk (Meiji) and subjected to primary antibody for 2 h and HRP-conjugated secondary antibody (GE Healthcare) for 30 min. Chemiluminescence was generated by ImmunoStar detection kit (Wako) or Immobilon (Millipore) and detected by LAS-3000 mini system (FUJIFILM).

### Cell-based *in vitro* Binding Assay

HEK293 cells were cultured in high glucose DMEM (Sigma-Aldrich) containing 10% FBS (HyClone), 50 unit/mL Penicillin, 50 mg/mL Streptomycin (Invitrogen), 2 mM L-Glutamine in 10% CO_2_ at 37°C. Cells were transfected with pDisplay encoding myc-tagged GluA1-GluA4-ATD or pCAGGS encoding FLAG-tagged Nrx or GFP with Lipofectamin 2000 (Invitrogen). On the following day, transfected cells were detached by phosphate-buffered saline (PBS) containing 5 mM EDTA and seeded on 12-mm coverslip coated with poly-L-lysine (PLL; Sigma-Aldrich) at 2 x 10^4^ cells/well. One hour after the seeding, cells were treated with vehicle, recombinant Cbln1-HIS or CPTX-HIS (23.6 nM, the final concentration as hexamer) for 4 h to overnight. For *in vitro* tripartite binding assay (fig. S3C), cells were further incubated with the conditioned medium containing Nrx1β(+4)-hFc for 4 h. Cells were fixed with 4% PFA/PBS for 15 min and washed with PBS three times. After blocking with 3%BSA (Sigma-Aldrich) /PBS for 30 min without permeabilization, cells were further incubated with primary antibody against the tag (HIS, HA or hFc) attached to the ligand in 3% BSA/PBS for 2 h at room temperature (RT) or 24 h at 4 °C. Following three washes with PBS, cells were permeabilized with PBS containing 0.1% Triton-X 100 (Sigma-Aldrich) and 3% BSA. Cells were stained with 1^st^ antibodies against tag (FLAG, Myc) of the binding receptor for 2 h at RT or 24 h at 4°C, followed by PBS wash and respective 2^nd^ antibodies for 30 min. Following the PBS wash, coverslips were mounted on a slide glass with Fluoromount-G. Samples in which HEK cells expressed GFP were not stained by 1^st^ antibodies against GFP but treated with the same 1^st^ antibodies against the tag of the receptor (FLAG or Myc) and the auto-fluorescence of GFP protein were detected by microscopy.

### *in vitro* Synapse Formation Assay

#### Hippocampal culture.

Preparation from embryonic day 16-17 (E16-17) wild-type ICR mice was performed as described with some modification (*32*). Neurons were plated at a density of 2 x 10^5^ cells on 12-mm cover glass coated with PLL and maintained in Neurobasal cell culture medium (Invitrogen) containing NS21 supplement (*33*), 50 U/mL Penicillin, 50 mg/mL Streptomycin (Invitrogen), 2 mM L-Glutamine and 2% FBS (HyClone) in 5% CO_2_ at 37°C. One to two hours after the initial incubation, the culture medium was changed to the fresh medium without FBS. For presynapse induction by HEK cells (fig. S4A), neurons were co-cultured with 2 x 10^4^ HEK cells expressing GFP or Myc-tagged ATD of either GluA1–4 or GluD2 at day-in-vitro 4 or 5 (DIV4 or 5), 1 h later followed by the treatment with vehicle, recombinant Cbln1-HIS or CPTX-HIS (23.6 nM, the final concentration as hexamer) for 2 d. For AMPA receptor induction by HEK cells (fig. S5A), neurons were co-cultured with HEK cells expressing GFP or Nrx-3xFLAG and/or CPTX-HIS at DIV9 for 2 d. For synapse induction by beads, neurons were treated with Dynabeads (M-280, streptavidin) coated with Cbln1-HIS or CPTX-HIS through anti-HIS biotin (MBL) for 1–2 d (DIV4–6 for VgluT1 induction in fig. S4, DIV20–22 for Syp, Nrx and GluA1–3 in Fig. 2BC and DIV30–32 for GluA4 induction in fig. S5B). To label axon and dendrite, GFP was transfected with Lipofectamine 2000 at DIV6. Following the incubation, the cells were fixed with 4% paraformaldehyde in PBS for 5 min at room temperature. Cells were washed with PBS and fixed with chilled MeOH for 8 min on ice, followed by the wash with PBS. Cells were permeabilized and blocked with PBS containing 0.1 % Triton-X 100, 3% BSA and stained with primary antibodies diluted in blocking buffer for 2 h at RT or 24 h at 4°C, followed by PBS wash and secondary antibodies diluted in blocking buffer for 30 min. Following PBS wash, coverslips were mounted on a slide glass with Fluoromount-G (Southern Biotech).

#### Cerebellar culture

Cerebellar cultures were prepared from postnatal day 0–3 (P0–3) *Cbln1*-null mice (on ICR background) with minor modifications (*32*). Neurons were plated at a density of 2 x 10^5^ cells on 12 mm cover glass coated with PLL and were maintained in DMEM/F12 (Sigma-Aldrich) containing 100 μM putrescine, 30 nM sodium selenite, 0.5 g/mL tri-iodothyronine, 0.25 mg/mL bovine serum albumin, 3.9 mM glutamate and N3 supplement (100 μg/mL apo-transferrin, 10 μg/mL insulin and 20 nM progesterone) and 10% FBS in 5% CO_2_ at 37°C. One to two hours after the initial incubation, the culture medium was changed to the fresh medium without FBS. Co-culturing with HEK cells (at DIV4 or 5 for 2 days), fixation, permeabilization and staining were performed as described for the hippocampal culture.

#### Quantification of Signal Intensity in *in vitro* Assay

Fluorescence images of cultured samples were captured using a CCD camera attached to a conventional fluorescence microscope (BX63, Olympus) or a confocal laser microscope (FV1000, Olympus) by 40x objective for the detection of HEK cells or by 60x objective with x5 digital zoom for the detection of beads. The same parameters, such as exposure time and excitation power, were used for the sample set to be compared, except for GFP signals. Analyses were performed semi-automatically by the customized macro using Image J (Fiji). Background subtraction using the rolling ball method (20–100 px) was applied to images to reduce the background noise. Regions of interest (ROIs) of HEK cells expressing receptors or beads were determined by intensity and size which fulfill the threshold (Otsu or Triangle) and particle definition. For quantification of signals on HEK cells, the signal intensity within the ROI was determined by subtracting the mean signal intensity outside ROI. For quantification of signals on beads, to minimize non-specific signals associated with the cross-reactivity of beads with antibodies, the signal intensity was determined by subtracting the mean intensity of beads located in the field where no cells were present (Fig. 2, B and C; fig. S4C). For fig. S5B, ROIs of beads were manually classified into two groups, those contacting and non-contacting with PV^+^ neurons, which were identified by PV and GluA4 immunoreactivities. The signal intensity was calculated by subtracting the intensity on non-contacting beads from that on contacting beads.

#### Immunocytochemistry and Immunohistochemistry

##### Detection of CPTX in hippocampal culture.

Dissociated hippocampal neurons were prepared from E17 embryos and cultured for 21 DIV. They were incubated overnight (between DIV20 and 21) with 1 μl of biotinylated CPTX (1.6 mg/mL, stock concentration), added directly to the culture medium. Neurons were fixed with 4% PFA, permeabilized with 0.1% Triton-X 100 in PBS for 10 min, washed three times and blocked (0.1% Glycine + 0.1% Tween-20 + 10% normal goat serum in PBS for Fig. 2D or 3% BSA + 0.1% Triton-X 100 in PBS for Fig. 3E) for 45 min at room temperature. Cells were incubated overnight with primary antibodies diluted in the blocking buffer and washed with PBS for three times. Cells were incubated with respective secondary antibodies for 1 h at room temperature. Coverslips were washed with PBS, incubated with or without DAPI (Invitrogen, reference number-D1306) for 10 min and mounted using Fluoromount aqueous mounting media (Sigma-F4680) or Fluoromount-G (Southern Biotech). For the acquisition of the images, complete Z stack spanning primary hippocampal cultures was acquired on a Leica TCS SP8-3X microscope equipped with a 405 nm Diode Laser and a white light laser (WLL) for excitation. The format for all images was set to 1024 x 1024 pixels with an optical zoom of 2 resulting in a pixel size of 90 nm in an X-Y plane. Images were taken with 400 lines/s and line averaging of 2. To improve image quality for representation, raw confocal images were deconvoluted using the Huygens Professional (SVI, v 15.10). The theoretical point spread function (PSF) parameters were automatically acquired from confocal lif-files. Within the deconvolution wizard, images were subjected to a manual background correction and signal-to-noise ratio optimization. The Optimized iteration mode of the GMLE was applied until it reached a Quality threshold of 0.01. For the representation of the images, the contrast was enhanced for the complete image by linear methods using Fiji for each channel separately.

##### Hippocampal sections.

For fig. S8C, the hippocampal slices used for *ex vivo* electrophysiological recordings from mice injected with CPTX and vehicle were fixed with 4% PFA for 30 min at room temperature. Cryoprotection was done in 30% sucrose overnight. Slices were further cut into 50μm thick sections. Membrane permeabilization was performed with 0.5% Triton-X 100 in PBS for 30 min. Sections were blocked with blocking buffer (0.1% Glycine + 0.1% Tween-20 + 10% Normal Goat Serum in PBS) for 90 min. Above mentioned antibodies were used for the detection of Bassoon (dilution 1:500) and CPTX-HIS (1:500), diluted in blocking buffer and applied for 2 d at 4°C. Sections were washed three washes with PBS followed by respective secondary antibodies (1:1000) for 3 h at room temperature. Sections were mounted using Fluoromount aqueous mounting media (Sigma-F4680). Complete Z stack spanning 50 μm thin hippocampal slices were acquired on a Leica TCS SP8-3X microscope. For acquiring images, the Leica objective HC PL APO CS2 63x/1.40 oil was used. Bassoon (405 nm), Thy1-GFP (488 nm), CPTX (546 nm) images were acquired with the confocal scanning mode. The format for all images was set to 512 × 512 pixels with an optical zoom of 3.04 resulting in a pixel size of 119 nm in an X-Y plane. Images were taken with 400 lines/s and line averaging of 4. For Fig. 4A; fig. S8, A and B; fig. S9; fig. S10; fig. S11), one day after the unilateral hippocampal injection of CPTX, mice were transcardially perfused with 4% PFA in 0.1 M sodium phosphate buffer (PB; pH 7.2) for 10 min under deep pentobarbital anesthesia, followed by post-fixation of dissected brain with 4% PFA in PB for 2 h at 4°C. Free-floating sections (50 μm thickness) were prepared by a microslicer (DTK-1000, Dosaka EM) and treated with 1 mg/mL pepsin (DAKO) in PBS containing 0.1% Triton X-100 and 0.2 N HCl (Wako) for 10 min at 37°C. For the high-resolution imaging of synaptic proteins, the brain was freshly frozen with liquid nitrogen after decapitation with isoflurane (Wako). Sections (10–20 μm thickness) were cut by cryostat (Leica), mounted on slide glass or coverslips and postfixed with 4% PFA in PB or 3% glyoxal (Sigma) for 2 h at RT. Following the wash with PBS containing 0.1% Triton X-100, sections were subsequently treated with 10% donkey serum for 30 min at room temperature, a mixture of primary antibodies overnight and a mixture of respective secondary antibodies. Finally, the sections were attached to slide glass and mounted with Vectashield (Vector) or prolong glass (Thermo). Fluorescence images were captured using confocal microscopies (SD-OSR, Olympus; LSM880, ZEISS) with the same parameters such as objectives, zoom, laser power, exposure time, gain and offset for the comparison of samples. The super-resolution images were obtained by the structural illumination with SD-OSR (Objective: 100x/1.49 oil with 3x zoom lens, pixel size: 40 nm/px, configuration: XY < 120 nm, Z < 300 nm) or by Airyscan with LSM880 (63x/1.40 oil with 3x digital zoom, 35 nm/px, XY < 120 nm, Z < 350 nm). Images obtained by Airyscan were 3D-reconstructed by Imaris (Bitplane). For the quantification of AMPAR clustering (fig. S9), images were obtained around the area of the injected side of the hippocampus (*Str. radiatum*) and the corresponding area of the contralateral uninjected side with 100x objective lens. Injected areas were confirmed by the HIS immunoreactivity. The dynamic range of GluA signals was first determined by maximal and minimal signal intensities in pair images of the injected and the contralateral area and then each signal intensity was normalized this value (max-min). Normalized GluA intensity histograms were calculated by dividing the normalized signal intensity in 40 bins at a logarithm scale. Thus, the signal intensity in the *n*^th^ bin equals 10^(*n* x (max-min) / 40)^. To analyze the bright pixels, the mean intensity + 3 x SD of the contralateral area was used as the threshold. Selected pixels were analyzed as particles by “Analyze Particles” command of image J without any limitation (all size and circularity), leading to the calculation of particle size, number, area fraction and intensity.

##### Cerebellar sections.

One day after the cerebellar injection of vehicle or CPTX into *Cbln1*-null mice, they were transcardially perfused with 4% PFA in 0.1 M sodium phosphate buffer (PB; pH 7.2) for 10 min under deep pentobarbital anesthesia, followed by post-fixation of dissected brain samples with 4% PFA in PB for 2 h at 4°C. Sections were prepared with a microslicer (DTK-1000, Dosaka EM) for free-floating sections (50 μm thickness). Sections were first treated with 1 mg/mL pepsin (DAKO) in PBS containing 0.1% Triton X-100 and 0.2 N HCl (Wako) for 10 min at 37°C. Following the wash of the sections with wash buffer (PBS containing 0.1% Triton X-100), sections were subsequently treated with 10% donkey serum for 30 min at room temperature, a mixture of primary antibodies diluted in wash buffer overnight, three washes, a mixture of respective secondary antibodies in wash buffer and three washes. Finally, the sections were attached to a slide glass and mounted with VECTASHIELD (Vector). Fluorescence images were captured using confocal microscopy (FV-1000, Olympus) at the same parameters such as objectives, zoom, laser power, gain and offset for the comparison of samples. Colocalization of signals was analyzed by the built-in plugin “Coloc 2” in Image J with the Fiji package (*34*).

##### Cortical sections.

One-month-old mice, to which miR vectors were transfected by *in utero* electroporation, were transcardially perfused with 3% glyoxal for 10 min under deep pentobarbital anesthesia, followed by post-fixation of the dissected brain with 3% glyoxal for 2 h at 4°C. Sections were prepared with a microslicer (DTK-1000, Dosaka EM) for free-floating sections (100 μm thickness). Following the wash of the sections with wash buffer (PBS containing 0.1% Triton X-100), sections were subsequently treated with 10% donkey serum for 30 min at room temperature, a mixture of primary antibodies diluted in wash buffer overnight, three washes, a mixture of respective secondary antibodies in wash buffer and three washes. Finally, the sections were attached to a slide glass and mounted with Vectashield (Vector). Z-stacked fluorescence images of NPs were obtained throughout the brain slice in the transfected cortical area at every 2 μm using a confocal microscope (SD-OSR, Olympus) by 20x objectives. ROI of soma area in layer 2/3 where the abundant expression of NPs was found in a large population of neurons was manually selected in an unbiased manner according to the GFP signal. For the quantification of the intensity of NPs in ROI by Image J, broad background signals were subtracted using the rolling ball method (50 px) and the signals in ROI was normalized by the surrounding signals enlarged at 5 px because the absolute signal intensity was strongly affected by the z-position of ROI in the brain section at 100 μm thickness.

##### Spinal cord sections.

Two to five days after the injection the mice were transcardially perfused with 3% glyoxal for 10 min under deep pentobarbital anesthesia, followed by post-fixation of the dissected spinal cord with 3% glyoxal overnight at 4°C and by cryoprotection in 30% sucrose/PBS for several days to be sliced. Spinal cords embedded in Tissue-Tek® O.C.T. compound (Sakura Finetek, #4583) were horizontally or coronally sectioned by cryostat (Leica) at 20-40 μm thickness in 4-5 mm around the epicenter of injury and the slices were mounted on the APS coated slide glass (Matsunami) at 200-400 μm interval. Following the wash with PBS containing 0.1% Triton X-100, sections were subsequently treated with 10% donkey serum for 30 min at room temperature, a mixture of primary antibodies overnight and a mixture of respective secondary antibodies. Finally, the sections were attached to slide glass and mounted with Fluoromount-G (Southern Biotech). Fluorescence images of all sections on the slide glass were captured as a virtual slide using conventional microscopy (BX63, Olympus) at 4x objective to determine the section at 1.4-1.6 mm upstream of the precise epicenter of the injury. The fluorescence image of the section was subsequently captured by confocal microscopies (SD-OSR, Olympus; 63x objective) around the ventral root in gray matter with the same parameters such as objectives, laser power, exposure time, gain and offset for the comparison of samples. The super-resolution images were obtained by Airyscan2 with LSM980 (63x/1.40 oil with 2.5x digital zoom, 35 nm/px, XY < 120 nm, Z < 350 nm). To define the particles of GluA4, VGluT2 and these intersections, the fluorescent images were subjective to the background subtraction (50 px), Laplace filter (9 x 9), box filter (5 x 5), auto thresholding (Otsu method), particle extraction (> 5 px), binarization and segmentation by watershed. The defined particles were used as ROI to quantify the size and mean intensity of each channel. Percentage of VGluT2 puncta with GluA4 was calculated by the number of VGluT2 particles accompanied by more than 1 px of defined GluA4 particles in the ROI divided by the number of all VGluT2 particles. All image processing was performed by ImageJ.

##### Golgi-Cox Staining and Dendritic Spine Analysis

Mice were killed and then decapitated. Brains were quickly removed from the skull and washed with bi-distilled water to remove blood from the surface. For the Golgi-Cox impregnation of neurons, the FD Rapid GolgiStainTM kit (FD NeuroTechnologies, #PK401) was used. Dye-impregnated brains were rapidly frozen on methyl butane and dry ice, embedded in Tissue-Tek® O.C.T. compound (Sakura Finetek, #4583), cryo-sectioned coronally at 100 µm thickness and directly mounted on gelatin-coated slides (FD NeuroTechnologies, #PO101) with the help of solution C provided in the kit. Sections were stained according to the manufactureŕs protocol and mounted with the embedding media Roti-Histokitt (Carl Roth, #6638.1). Secondary apical dendrites from CA1 pyramidal neurons with cell bodies located in the dorsal part of the hippocampus were imaged for spine analysis. Sections were imaged by an experimenter blinded to experimental conditions using a Leica Leitz DMRXE microscope as z-stacks with 0.25 µm interval between sections at a magnification of 100x. Between 7 and 8 dendrites of 60-80 μm length were imaged for each animal. The analysis was done by an experimenter blinded to experimental conditions using the Neurolucida software version 11 (MBF–Bioscience).

##### Electron Microscopy

For the *in vivo* Cbln1 and CPTX administration experiments, *Cbln1*-null or *GluD2*-null (C57/BL6J background, 3-6 months old) were perfused transcardially with 2% PFA, 2% glutaraldehyde in 0.1M PB (pH 7.2) under deep pentobarbital anesthesia 3 d after the injection into the cerebellum. Parasagittal microslicer sections of the cerebellum (300 μm) were postfixed for 2 h with 1% OsO_4_ in 0.1 M PB. After block staining in 1% aqueous uranyl acetate solution and dehydration with graded alcohols, the sections were embedded in Epon 812. Serial ultrathin sections (70 nm) were made using an ultramicrotome and stained with 2% uranyl acetate for 5 min and mixed lead solution for 2 min. Electron micrographs of the molecular layer were taken using an H-7100 electron microscope at a magnification of x4,000 and printed at a magnification of x16,000. For the quantitative analysis of PF–Purkinje cell synapses, the asymmetrical postsynaptic density in the cerebellar cortex was randomly photographed and the frequency of contacted and naked/free synapses was analyzed by using MetaMorph software as described previously (*35*). Three series of serial sections, each of which consisted of ten electron micrographs, were analyzed (*n* = 1-2 mice).

##### *In vitro* Electrophysiology

*Hippocampus (Figs. 4, S13 and S14).* All experiments were performed in 4- to 6-week-old C57BL/6J wild type, 4- to 6-week-old PV-*Cre*/tdTomato-FLEX double transgenic mice (for mEPSCs recordings from CA1 PV-positive interneurons), or 5-month-old 5xFAD mice (for mEPSCs recordings), or 15- to 18-month-old 5xFAD transgenic mice (for LTP recordings) as described previously (*36*). Animals were killed and then decapitated. Hippocampi from both hemispheres were isolated and placed on agar block to cut transverse slices (350 μm) with the vibrating microtome (VT1200S, Leica) in ice-cold solution containing (in mM): 2 KCl, 1 MgCl_2_, 2 MgSO_4_, 1.25 NaH_2_PO_4_, 26 NaHCO_3_, 1 CaCl_2_, 10 D-glucose, and 230 sucrose or 93 NMDG, 2.5 KCl, 1.2 NaH_2_PO_4_, 30 NaHCO_3_, 0.5 CaCl_2_, 10 MgSO_4_, 20 HEPES, 25 glucose, 5 sodium ascorbate, 3 sodium pyruvate (pH 7.4 with HCl). All solutions were saturated with 95% O_2_ and 5% CO_2_ and the osmolality was adjusted to 300 ± 5 mOsm. For CPTX *in vitro* treatments, slices were incubated in a 2-ml chamber for 4 h at room temperature in the ACSF solution containing (in mM): 124 NaCl, 2.5 KCl, 1.3 MgSO_4_, 1 NaH_2_PO_4_, 26.2 NaHCO_3_, 2.5 CaCl_2_, and 11 D-glucose. CPTX was added at a concentration of 20 μg/mL to slices in the chamber, whereas vehicle slices were incubated in ACSF without CPTX. Next, the slices were transferred to the recording chamber and were continuously perfused with the same ACSF for miniature excitatory postsynaptic currents (mEPSC) measurements. Picrotoxin (50 μM, Tocris) CGP 55845 (3 μM, Tocris), and tetrodotoxin (Tocris) were added to ACSF to block GABA_A_, GABA_B_ receptor and Na^+^ channels, respectively, and hence to isolate mEPSCs. Whole-cell patch recordings from CA1 pyramidal neurons were obtained using glass electrodes (4–6 MΩ, outer diameter of 1.5 mm; wall thickness of 0.315 mm; Hilgenberg, Germany) with a resistance of 4–6 MΩ. The glass electrodes were filled with a solution containing (in mM) 140 K-gluconate, 8 NaCl, 0.2 CaCl_2_, 10 HEPES, 5 QX314Br, 0.5 Na_2_GTP, and 2 Mg_2_ATP (pH adjusted to 7.2 with KOH and osmolality adjusted to 290 mOsm). The membrane potential was clamped at –70 mV. For mIPSC recording, NBQX (25 μM, AMPAR antagonist, Tocris), D-AP5 (50 μM, NMDAR antagonist, Tocris), CGP 55845 were added into ACSF solution. The glass electrodes were filled with a solution containing (in mM): 120 CsCl, 8 NaCl, 0.2 MgCl_2_, 10 HEPES, 2 EGTA, 5 QX 314-Br, 0.3 Na_2_GTP, and 2 Mg_2_ATP (pH 7.2, 290 ± 3 mOsm).

ACSF solution containing (in mM): 120 NaCl, 2.5 KCl, 1.5 MgCl_2_, 1.25 NaH_2_PO_4_, 24 NaHCO_3_, 2 CaCl_2_, and 25 D-glucose was used for field excitatory postsynaptic potential (fEPSP) recording. Thin glass electrodes (outer diameter 1.5 mm, wall thickness 0.188 mm, ∼2 MΩ) filled with ACSF were used for stimulation and recording of fEPSPs. The Shaffer collaterals pathway was stimulated using a glass electrode and theta-burst stimulation (TBS) trains were applied three times to induce LTP (*37*). The stimulation intensity was determined based on the input-output curve and was set to give fEPSPs with the slope ≈50 % of the supramaximal fEPSP. Single stimuli were repeated every 20 s for at least 10 min for baseline recording before and for 60 min after LTP induction. The paired-pulse ratio (PPR) was evaluated at 50-ms intervals under the same conditions. mEPSC and mIPSC events were analyzed offline with Mini Analysis Program (6.0.3., Synaptosoft, USA).

Recordings were obtained using an EPC-10 amplifier (HEKA Electronik). The recordings were digitized at 10-20 kHz and filtered at 1-3 kHz. Values are expressed as means ± SEM. Statistical differences were determined using the t-test, one- and two-way ANOVA, and Holm-Sidak *post-hoc* test.

##### Hippocampus (fig. S15).

All experiments were performed in 6- to 9-week-old C57BL/6J mice. Coronal hippocampal slices (300-μm thick) were prepared by tissue slicers (7000-smz, Campden instruments,) 3 days after vehicle or CPTX injection and stored at 34°C in the ACSF. Whole-cell patch-clamp recordings were made from CA1 pyramidal neurons at 32–33°C. The resistance of the patch pipettes was ∼4 MΩ when filled with an intracellular solution composed of (in mM): 150 Cs-gluconate, 10 HEPES, 4 MgCl_2_, 4 Na_2_-ATP, 1 Na_2_-GTP, 0.4 EGTA, 5 QX-314, pH 7.25 (adjusted with CsOH), 292 mOsm/kg) for measurements of the input-output relationship and the paired-pulse ratio of temproaromic (TA) pathway to the *lacunosum moleculere* (Lm) amplitudes. The solution used for the slice storage and recording consisted of the following (in mM): 125 NaCl, 2.5 KCl, 2 CaCl_2_, 1 MgCl_2_, 1.25 NaH2PO4, 25 NaHCO_3_, 25 D-glucose with 0.1 Picrotoxin, bubbled continuously with a mixture of 95% O_2_ and 5% CO_2_. To evoke the TA-Lm transmission, square pulses were applied at various stimulus intensities (20 μs, 50−600 μA) through a stimulating bipolar electrode made from theta tubing (Harvard apparatus, 30-0117) placed on Lm (∼50 μm medially away from the recording pipette). Holding potential was −70 mV (liquid junction potential was not corrected) and Rs was monitored during the experiment by applying −5 mV test pulse during recordings. Recordings with >20% Rs change were discarded. The paired-pulse ratio of EPSC amplitudes were recorded at various intervals (20-500 μs). Decay times were analyzed by the responses induced by stimulations at 600 μA. Current responses were recorded with a HEKA EPC-9/2 (HEKA) at a 20 kHz sampling rate with a low pass filter (2.9 kHz), and Igor Pro 7 (Wavemetrics) was used for data acquisition and analysis.

##### Cerebellum.

Parasagittal cerebellar slices (200-μm thick) were prepared from 4- to 6-week-old *GluD2*-null or *Cbln1*-null mice at 3 days after vehicle, Cbln1, NP1_PTX-3Cl_ or CPTX injection, as described previously (*2*). Whole-cell patch-clamp recordings were made from PCs using a 60× water-immersion objective attached to an upright microscope (BX51WI, Olympus) at room temperature. The resistance of the patch pipettes was 1.5−3 MΩ when filled with an intracellular solution composed of (in mM): 150 Cs-gluconate, 10 HEPES, 4 MgCl_2_, 4 Na_2_ATP, 1 Na_2_GTP, 0.4 EGTA, and 5 lidocaine *N*-ethyl bromide (QX-314) (pH 7.3, 298 mOsm/kg) for measurements of the input-output relationship and paired-pulse ratio of PF-EPSC amplitudes; 65 Cs-methanesulfonate, 65 K-gluconate, 20 HEPES, 10 KCl, 1 MgCl_2_, 4 Na_2_ATP, 1 Na_2_GTP, 5 sucrose, and 0.4 EGTA (pH 7.25, 295 mOsm/kg) for LTD recordings. The solution used for the slice storage and recording consisted of the following (in mM): 125 NaCl, 2.5 KCl, 2 CaCl_2_, 1 MgCl_2_, 1.25 NaH_2_PO_4_, 26 NaHCO_3_ and 10 D-glucose, bubbled continuously with a mixture of 95% O_2_ and 5% CO_2_. Picrotoxin (100 μM, Sigma) was always present in the saline to block the inhibitory inputs. To evoke the PF-EPSCs, square pulses were applied at various stimulus intensities (10 μs, 0−200 μA) through a stimulating electrode placed on the molecular layer (∼50 μm away from the pial surface). Selective stimulation of the PFs was confirmed by paired-pulse facilitation of EPSC amplitudes at a 50-ms inter-stimulus interval. Current responses were recorded with an Axopatch 200B amplifier (Molecular Devices), and pClamp software (version 9.2, Molecular Devices) was used for data acquisition and analysis. Signals were filtered at 1 kHz and digitized at 4 kHz.

##### CPTX Injection

###### Acute intrahippocampal injection.

We used a digitally controlled infusion system (UltraMicroPump, UMP3, and Micro4 Controller, WPI) fed with a 10 μl Hamilton syringe and a NanoFil (35 GA) beveled needle. The mouse was first anesthetized with 1-3 % isoflurane and put into the stereotaxic frame (Narishige, Japan). 1 μl CPTX (1.49 mg/mL) or vehicle (HBS buffer as used for elution of CPTX, composed of 10 mM HEPES pH 7.4, 150 mM NaCl) was injected at a rate of 3 nl/s bilaterally. We used the following coordinates for bilateral injection: AP = −2.0 mm from Bregma and L = ±1.5 mm; DV = 2.0 mm from the brain surface according to the mouse brain atlas(*38*).

###### Acute intracerebellar injection.

Mice were anesthetized with an intraperitoneal injection of ketamine/xylazine (80/20 mg/kg body weight; Daiichi-Sankyo/Sigma). As described previously (*35*), a small hole in the occipital bone was made with a dental drill, and the dura matter was ablated. A glass needle was inserted into the vermis of lobule VI/VII at ±0.7 mm from the midline at the depth of around 200 μm. Approximately 7 μL of vehicle, Cbln1-HIS or CPTX-HIS (7.7 μM) was bilaterally injected at a rate of 10-40 μl/h.

###### Acute intrahippocampal injection through guide cannulas.

Intrahippocampal guide cannula was implanted as described previously (*39–41*) with minor changes. In brief, mice were anesthetized with 1–3% isoflurane (Baxter International Inc.) mixed with O_2_ through a vaporizer (Matrx VIP 3000, Midmark). Prior to any surgical manipulation, the mouse was given the analgesic carpofen (5 mg/kg b.w. s.c., Rimadyl, Pfizer Pharma GmbH) and the skin was cleaned by 75% ethanol, followed by 10% povidone iodine (Dynarex) and an additional analgesic xylocaine (a pumpspray, 10 mg lidocaine, Astra Zeneca GmbH). The mouse was placed in a stereotaxic frame (Narishige), and all next procedures were performed under a surgical binocular microscope (Labomed Prima DNT, Labo America Inc.) and on a heating pad (DC Temperature Controller, WPI) to maintain mouse body temperature constant (34–36 °C). The mouse scalp skin was circular incised (ϕ 10 mm) and removed. The edges of the skin were processed with 75% ethanol and xylocaine. The scalp bone was carefully cleaned with 75% ethanol and 3% H_2_O_2_ and dried under 40 °C (Steinel GmbH, HG-2310 LCD). After marking coordinates for implantation, 4 small holes for anchoring screws were drilled in the frontal and parietal bones by using a dental micro motor (Eickemeyer). Then all four screws were covered with acrylic dental cement Paladur (Heraeus Kulzer GmbH) leaving the marked areas for cannulas free of acrylic. Implantation began with drilling of a hole for the left cannula. The cannula was stereotaxically implanted by using a self-made universal holder for cannulas (small curved tweezers from FST with a metal ring). During implantation, the cannula gently touched the surface of the brain and only then it was secured with a small amount of acrylic. Next, the right cannula was similarly implanted and secured with acrylic. Coordinates for bilateral cannulas (AP: 2.0 mm; L: ±1.5 mm from Bregma, the bottom end of cannulas should touch the surface of the brain), were set according to the mouse brain atlas (*38*). After implantation of cannulas more dental cement was used to secure the whole system on the bone. After the surgery, which took for 60 min, mice were returned to the home cages. Carpofen (5 mg/kg b.w. s.c., Rimadyl, Pfizer Pharma GmbH) was used as a postoperative analgesic. The intrahippocampal injections and behavioral tests were performed after the mice had fully recovered at least 5–7 d after surgery. Injections were done using a 10 μl Hamilton syringe as described in *Acute intrahippocampal injection* section but being inserted in the guide cannula under brief sedation with 1% isoflurane.

*Acute intraspinal injection to SCI induced animals* 0.5 μl of CPTX (1 μg/μl), Cbln1(1 μg/μl), Chondroitinase ABC (0.5 U/μl; Sigma Aldrich, C2905) or vehicle (HBS buffer) was injected in the proximity of the site of injury by electrical micro-injector (BJ110, BEX CO., LTD.) through glass capillaries (3-000-203-G/X, Drummond Scientific Company). Following injection, the muscle layers and skins were again closed by sutures.

###### *In utero* Electroporation

*In utero* electroporation (IUE) was performed on E14.0 in the cortex of ICR mice as described previously (*42*). Briefly, plasmid DNA was injected into the third ventricle by a glass pipette and a set of electrical pulses was applied 5 times. Positive electrodes were placed onto the cortical side. Plasmid DNA was dissolved in 21 mM HEPES, 137 mM NaCl, 5 mM KCl, and 0.7 mM Na_2_HPO_4_ at a concentration of 1 mg/mL.

###### Surgical Procedure of Spinal Cord Injury (SCI)

Mice (9–11 weeks old) were subject to hemisection- or compression-induced SCI. After the anesthesia, the spinal cord was surgically exposed and the dorsal column at the 10^th^ thoracic vertebrae (T10) was dissected by micro-scissors and a scalpel. The muscle layers and skins were closed by sutures. Animals were recovered from the anesthetic by the administration of antagonist. Mice were subject to compression-induced damage with a commercially available SCI impactor device (Infinite Horizon Impactor; Precision Systems and Instrumentation, Lexington, NY) (*43*). We used a 70-kdyne impact force. We monitored the strength and the duration of the impact of compression through the automated recording system in the SCI impactor device.

###### Spatial navigation in a labyrinth.

To validate whether CPTX might have an effect on the hippocampus-dependent brain function, we used a labyrinth, 3D-printed dry maze developed at the DZNE. In short, before implantation of cannulas, mice were put on the light food restriction diet (LFRD) scheme, namely, *ad libitum* food was removed from mice and was given daily manually at the same day time with amount comparable with normal consumption (4-5 g per day per mouse). The weight of all mice was monitored daily and kept within a 10–15% reduction as maximum. Before the cannulation of mice, we ran labyrinth habituation and training for 2-3 weeks. Once all mice got familiar with the labyrinth, for the next experiments with CPTX, we changed the room to introduce new landmarks and cues, and we changed the configuration of the labyrinth to be more complex (Fig. 4I). Testing in the labyrinth began on day 3 (d3) after CPTX or Vehicle injection. A mouse was placed into a starting zone in the labyrinth and given 20 min to find a hidden reward as a food colorless and odorless round pellets (10 mg, AIN-76A, #1811213, TestDiet, USA). Two hours later the mouse was placed again in the same labyrinth with the same landmarks and cues in the room for 10 min to find the reward. Next day, d4, the mouse was placed again into the same labyrinth while the reward zone was changed, in a such manner that the mouse had to re-learn a new position of the reward within 10 min. Two hours later, in the retrieval session (5 min), the mouse had to find the new location of the reward.

The position of mice in the labyrinth was recorded on an HP workstation Z400, with Xeon computer (CPU 3.07 GHz, 6 GB RAM) using a USB video camera (Microsoft, USA) and special behavioral video acquisition and analysis software (AnyMaze, USA). All recorded movies were then analyzed using AnyMaze by a trained observer blind to the mouse genotype and treatments. AnyMaze was run in a tracking mode to trace coordinates of mice (10 frames/s) and compute distance traveled from the starting zone to the hidden reward zone. The criteria for successful performance was that during encoding and retrieval sessions a mouse finds the hidden reward (7-8 pellets) and eats all pellets. Travel distances in meters were computed as follows: a mouse must find the reward and take at least 1 pellet. This time point, as a beginning of the association of the reward location in the labyrinth and all landmarks and cues in the room, was used to calculate travel distances which were averaged and presented as a mean ± SEM.

###### Contextual fear conditioning.

To validate the action of CPTX in another hippocampus-dependent cognitive task but in the different codomain, we chose contextual fear conditioning (CFC). Shortly after spatial navigation in the labyrinth, at day 5 all mice were first recorded in a neutral context (context B, 5 min) and then 1 h later they were put into a conditioned context (context A, 5 min) and given 3x foot shocks, each with 0.5 mA, 1 s in duration and separated by 1 min. Two contexts, A and B, were different in terms of walls (chess-like black-and-white pattern versus grey color), smell (different cleaning solutions) and context A had a metal grid (to deliver foot shock), while context B had a white plastic floor. The next day after CFC, on day 6 after CPTX/Vehicle injection, for retrieval sessions, all mice were first put in the context A for 5 min, then at least 1 h later in the context B for 5 min. CFC memory in retrieval sessions was measured as % of the time a mouse was freezing in contexts A and B. Freezing of mice was defined as total immobility of animals (except for breathing) with a characteristic tense fearful posture. The analysis of data was done by using AnyMaze software. The fear conditioning system consisted of a touch-pad controller and a conditional sound-attenuated cabinet and chambers from Ugo Basile, Italy.

###### Rotor-rod test and gait analyses.

To examine the skilled motor coordination in the mice, the rotor-rod test was performed. As this phenotype is clearly observed on the ICR genetic background (*2, 32, 35*), we used in 4- to 6-week-old *Cbln1*-null mice (ICR background). The rotor-rod test was performed 1 d before and three days after the injection. Six trials were performed at 20 rpm with a 30-s interval in each trial and the latency to fall from the rotor-rod was measured (maximum score, 120 s). For gait analyses, we used 4- to 6-month-old *Cbln1*-null and *GluD2*-null (C57/BL/6J) mice. In addition, 4- to 6-week-old *Cbln1*-null mice (ICR background) were used. Mice were habituated to the experimenter and the behavior room from 2-3 d before the test. Then, the mice were habituated to the gait analysis apparatus made of transparent Plexiglass (6 cm x 85 cm) and trained to walk straight forward heading their home cage. Each step during the walking was recorded by a video camera across the transparent floor at 30 frames per second. The time point and location of the hind limb to attach and detach on the floor were manually determined in a blind manner with the program made by Hot Soup Processor. These analyses were performed more than 4 times for each mouse 1 d before and 3 d after the injection to calculate four gait parameters as follows: i) The coordinated step ratio was calculated as the number of alternate left/right steps divided by the number of total steps; ii) The stride length of each step along the X-axis (plus direction to home cage); iii) the stride speed, and iv) stride irregularity (the standard deviation of stride length during the single journey divided by the average stride length).

###### Locomotor recovery after SCI.

The locomotor recovery assessment was performed using video recording as previously reported (*43*). BMS (Basso Mouse Scale) open-field scoring and footfall tests were performed weekly during the 6–8 weeks following SCI. The BMS score was evaluated by at least two independent investigators who were unaware of the experimental groups. Mice were excluded if they had an incomplete injury (BMS score > 0) on the 2^nd^ day after SCI. To quantify the recovery effect, the increase in BMS score between the 1^st^ and 2^nd^ week after SCI was calculated as ΔBMS/week = BMS score (2^nd^ week) - BMS score (1^st^ week).

For footfall tests, a mouse was placed on a wire-mesh grid and videotaped for 5 min while on the grid. Mice that walked longer than 3 min with more than 70 steps were subjected to scoring by 3 independent examiners. The total number of footfalls from the bars during the total walking time was counted.

---

**Movie S1. Gait performance before and after the injection of CPTX in *Cbln1*-null mice on the C57BL/6 background.** Representative movie of gait and footprint of *Cbln1*-null mice (C57BL/6 background) before (top) or 3 days after (bottom) the injection of CPTX. Related to fig. S6C.

**Movie S2. Gait performance after the injection of CPTX in *Cbln1*-null mice on the ICR background.** Representative movie of gait and footprint of *Cbln1*-null mice (ICR background) 3 days after the injection of mock (vehicle injection) (top), Cbln1 (middle) or CPTX (bottom). Related to fig. S7A.

**Movie S3. Gait performance before and after the injection of CPTX in *GluD2*-null mice on the C57BL/6 background.** Representative movie of gait and footprint of *GluD2*-null mice (C57BL/6 background) before (top) or 3 days after (bottom) the injection of CPTX. Related to **fig. 3E**.

**Movie S4. Effects of acute CPTX injections on motor performance 4 weeks after SCI. Representative movie of free walking at 4 weeks after SCI.** The mouse which received the CPTX injection immediately after SCI has a mark on the tail. Related to **Fig. 5G**

**Movie S5. Effects of acute CPTX injections on footfall test 6 weeks after SCI.** Representative movie of footfall test at 6 weeks after SCI. CPTX was injected immediately after SCI. Related to figure S15D.

**Movie S6. Effects of the mock injection at 1week on footfall test at 2 weeks after SCI.** Representative movie showing the effect of mock injections on footfall test. Related to figure S15C.

**Movie S7. Effects of the CPTX injection at 1 week on footfall test 2 weeks after SCI.** Representative movie showing the effect of CPTX injections on foot fall test. Related to figure S15C.

**fig. S1.**
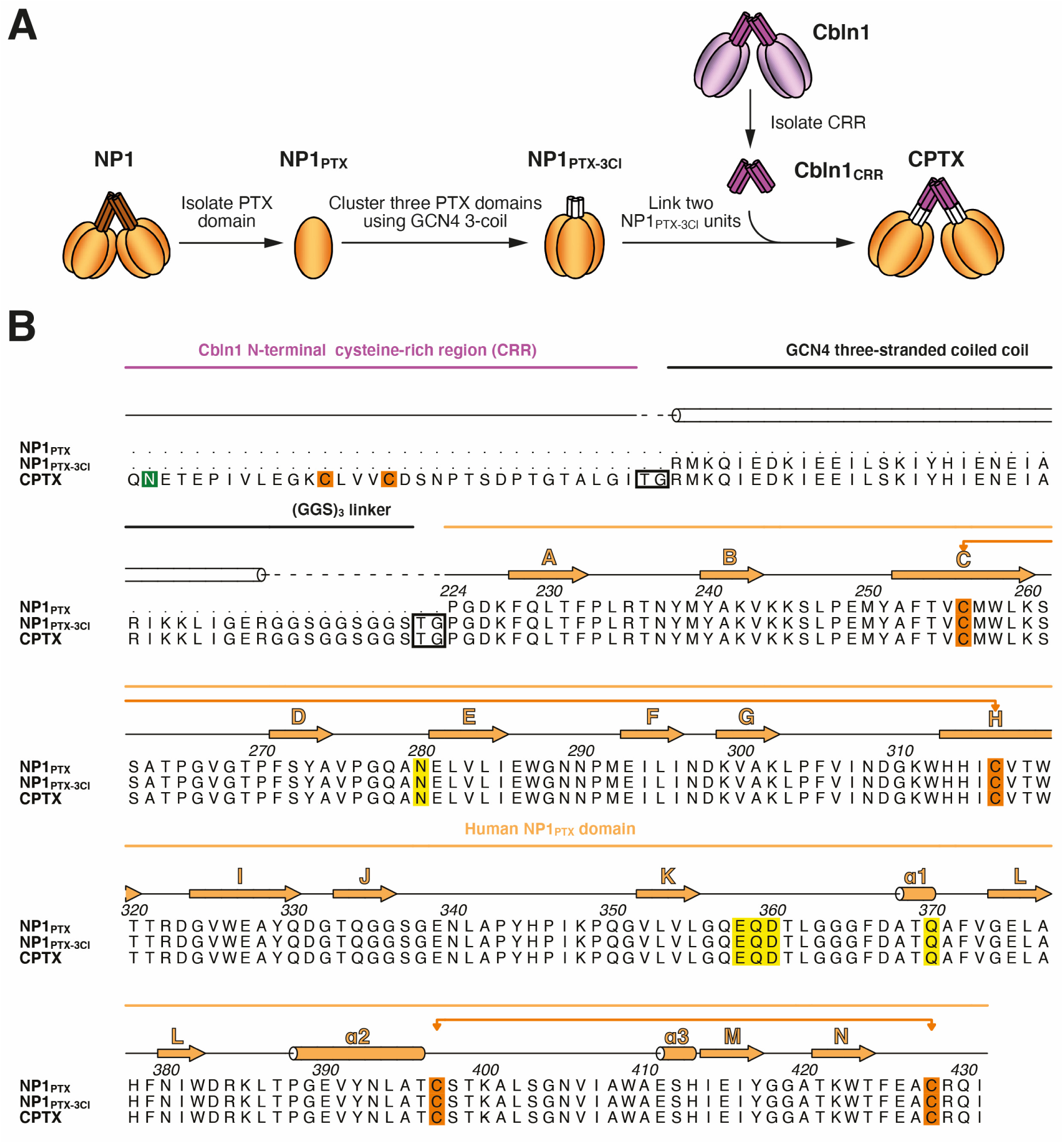
Sequence alignment of NP1_PTX_, NP1_PTX-3Cl_ and CPTX. **(A)** Construction of the synaptic organizer CPTX. **(B)** Sequence alignment of NP1_PTX_, NP1_PTX-3Cl_ and CPTX. The Cbln1 N-terminal cysteine-rich region (CRR), the GCN4 three-stranded coiled-coil, the (Gly-Gly-Ser)_3_ linkers, as well as the PTX domain and its secondary structure elements, are annotated above the alignment. Cysteine residues within the PTX domain that participate in disulfide bonds are shown in orange and connected with orange lines. Conserved calcium-coordinating residues are highlighted in yellow.

**fig. S2.**
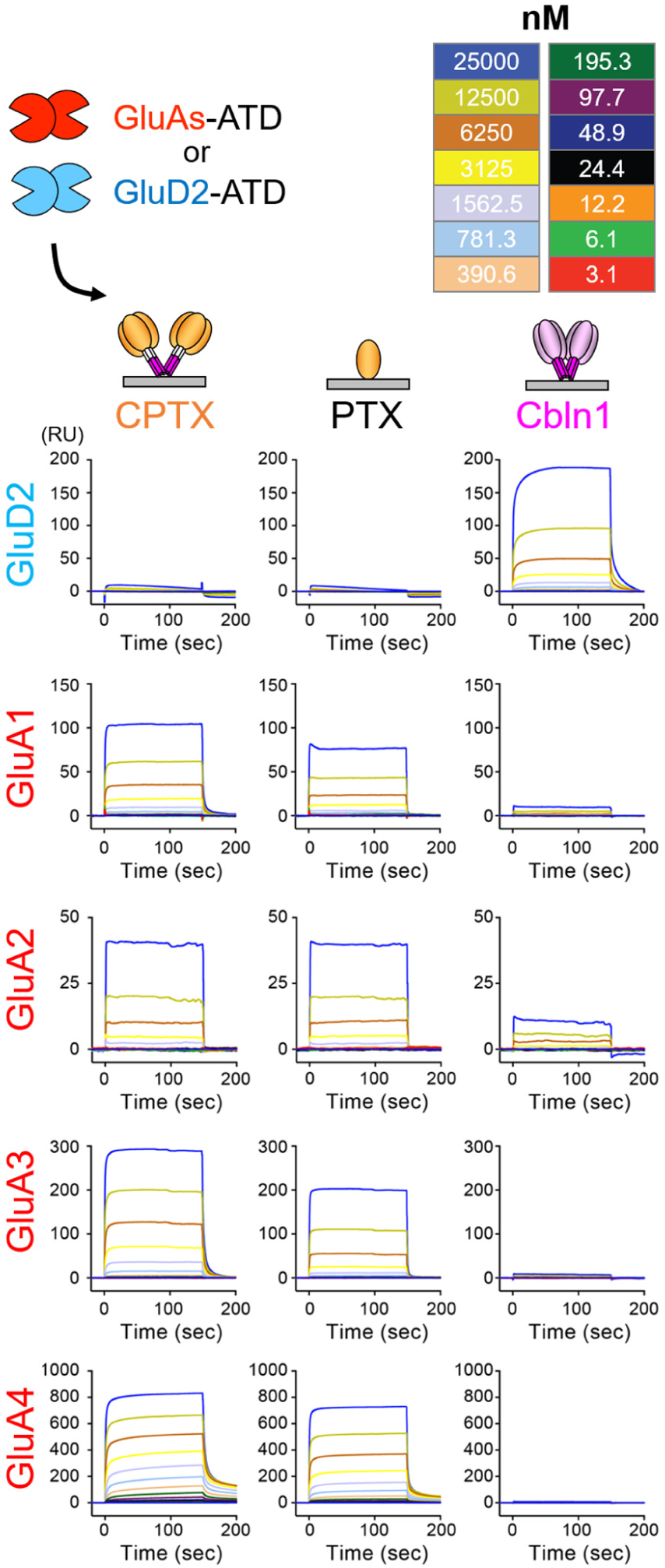
SPR measurements of the interactions between CPTX and the GluA1-4 and GluD2 amino-terminal domains. Schematic representation of the SPR setup, and sensorgrams for the interaction of Cbln1, NP1PTX and CPTX with mouse GluA1-4 ATD and GluD2 ATD.

**fig. S3.**
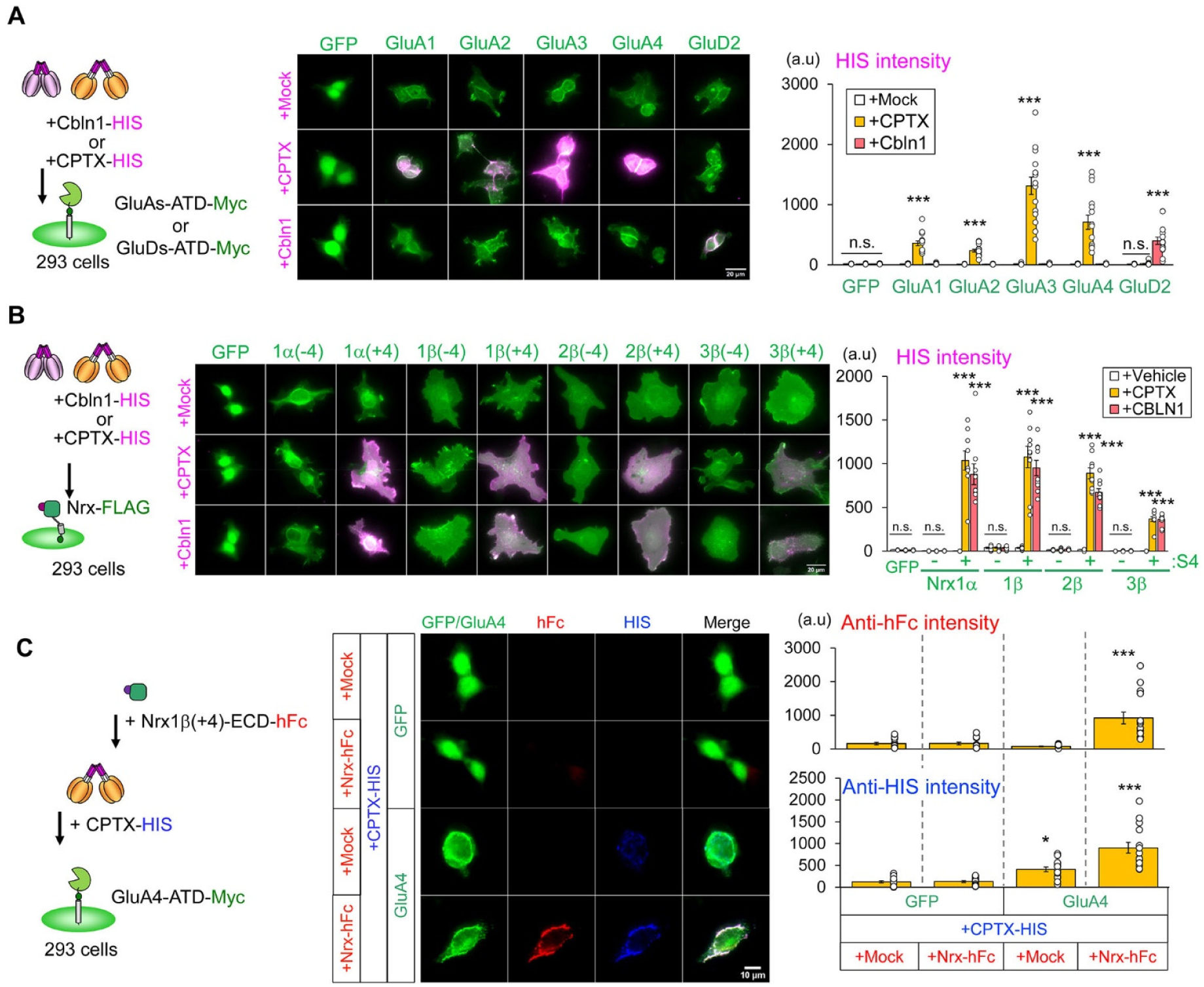
Cell-based binding assays show CPTX binds AMPARs and Nrx. (**A, B**) CPTX binds AMPARs and Nrx(+4) GluD2. Vehicle (mock treatment), HIS-tagged Cbln1 or CPTX-HIS (23.6 nM) was incubated with HEK cells expressing GFP or myc-tagged ATDs of GluA1–GluA4 AMPARs or the myc-tagged ATD of GluD2 (**A**). Cbln1-HIS or CPTX-HIS was incubated with HEK cells expressing Flag-tagged various Nrx isoforms with or without S4 or GFP (**B**). The intensities of HIS immunoreactivity (magenta) on green-positive cells were quantified in the right graphs. The bars represent mean ± SEM values. \*\*\**P* < 0.001, n.s. means not significant, *n* = 5–15 fields in more than 3 independent experiments, one-way ANOVA followed by Bonferroni test (compared with the other ligand treatment in each receptor). (**C**) CPTX mediates a tripartite complex consisting of Nrx(+4), CPTX and GluAs. CPTX-HIS (23.6 nM) was incubated with HEK cells expressing GFP or Myc-tagged GluA4-ATD, followed by the conditioned medium of HEK cells transfected with the human Fc (hFc)–tagged extracellular domain (ECD) of Nrx1β(+4) or empty vector (mock treatment). The intensities of hFc (red) and HIS (blue) immunoreactivity on Myc-positive cells (green) were quantified in the right graph. The bars represent mean ± SEM values. \**P* < 0.05, \*\*\**P* < 0.001, *n* = 10–15 fields in 3 independent experiments, one-way ANOVA followed by Tukey’s test (compared with GFP + CPTX-HIS + Nrx-hFc).

**fig. S4.**
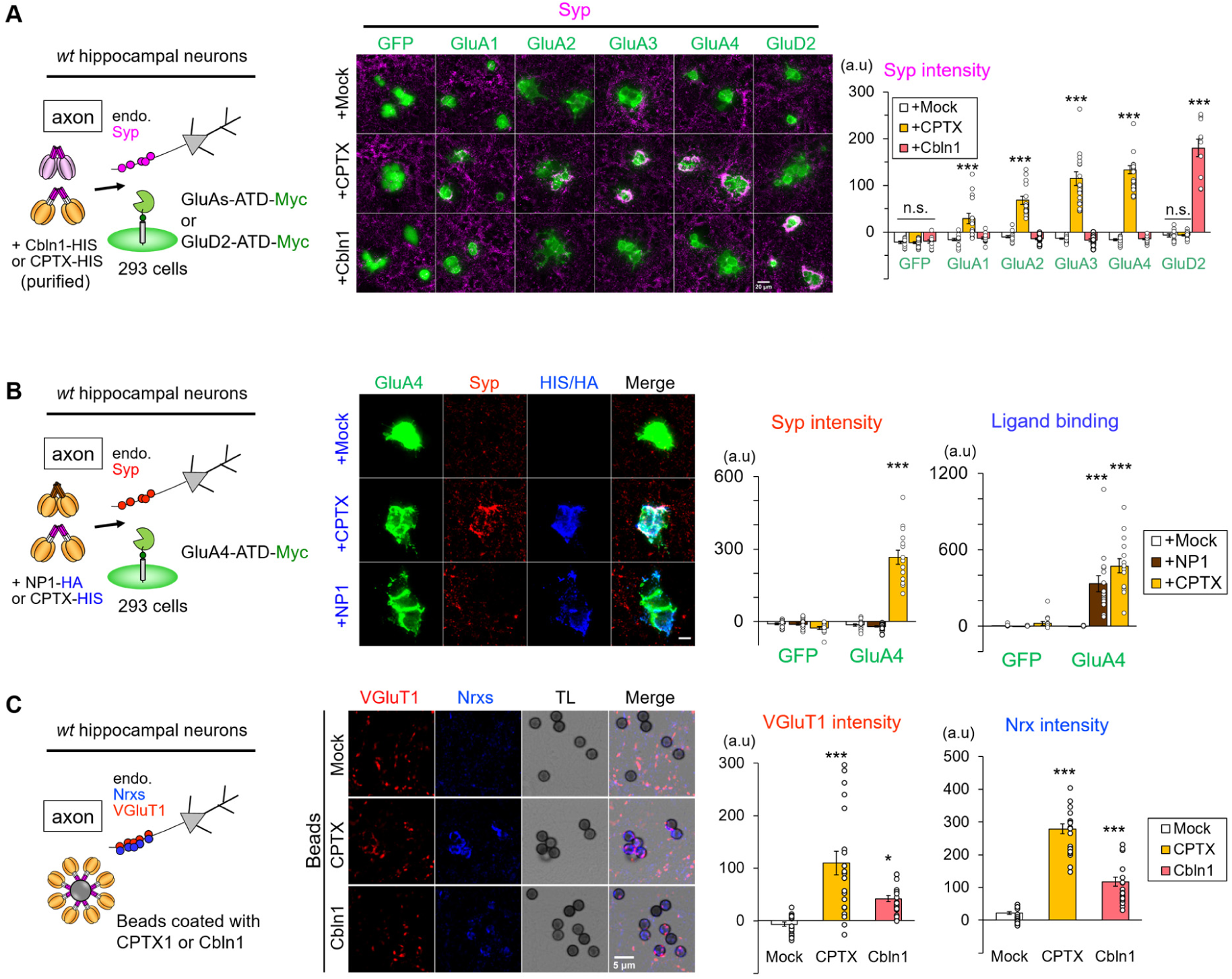
CPTX induces excitatory presynaptic sites in cultured hippocampal neurons. (**A**) Application of recombinant CPTX, but not Cbln1 or vehicle (mock treatment), induces accumulation of synaptophysin (Syp). Representative immunocytochemical images show accumulation of endogenous presynaptic sites (magenta; Syp, upper panels; Nrx, lower panels) of co-cultured hippocampal neurons on HEK cells displaying the ATDs of AMPA receptors, but not GluD2 (green). Scale bars, 30 μm. The right graph shows averaged intensities of Syp immunoreactivity on HEK cells expressing GluA or GluD2 ATDs. The bars represent mean ± SEM values. \*\*\**P* < 0.001, *n* = 10–15 fields in 4 independent experiments, Bonferroni test (compared with the other ligand treatment in each receptor). (**B**) Application of recombinant NP1 does not induce accumulation of Syp. HEK cells overexpressing GluA4-ATD were treated with the conditioned medium from HEK cell overexpressing empty vector (mock treatment), HA-tagged full-length NP1 or HIS-tagged CPTX (blue). Representative immunocytochemical images show that while both NP1 and CPTX (blue) bind to HEK cells expressing GluA4-ATD, only CPTX accumulates endogenous Syp (red). The right graph shows averaged intensities of Syp and HIS/HA immunoreactivity on HEK cells expressing GluA4-ATD. The bars represent mean ± SEM values. *n* = 15 fields in 3 independent experiments, one-way ANOVA followed by Tukey’s test (\**P* < 0.001 compared with the mock ligand treatment). (**C**) Beads coated with Cbln1 or CPTX, but not mock (beads treated with vehicle), accumulate endogenous Nrx (blue) and VGluT1 (red) immunoreactivities in axons of co-cultured wild-type hippocampal neurons. The right graph shows averaged intensities of VGluT1 and Nrx immunoreactivities on beads. \**P* < 0.05, \*\**P* < 0.001, *n* = 20 fields in 3 independent experiments, one-way ANOVA followed by Tukey’s test (compared with mock).

**fig. S5.**
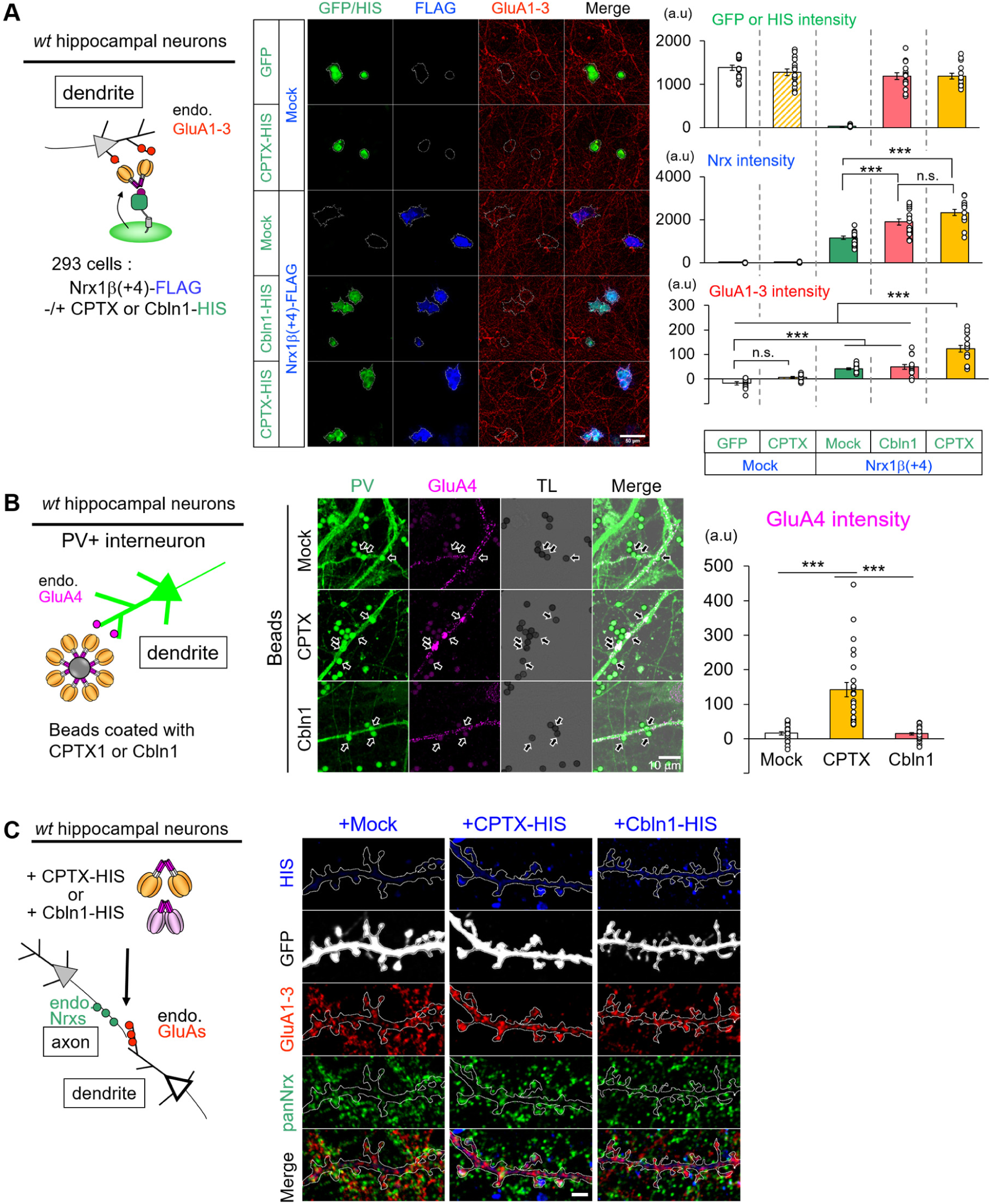
CPTX recruits endogenous postsynaptic AMPARs in hippocampal neurons. (**A**) Co-expression of CPTX and Nrx1β(+4) facilitates the accumulation of endogenous AMPARs in co-cultured hippocampal neurons. Hippocampal neurons were co-cultured with HEK cells expressing a combination of Nrx1β(+4) and HIS-tagged CPTX or Cbln1-HIS or empty vector (mock). Representative immunocytochemical staining shows that endogenous GluA1–3 (red) in hippocampal neurons accumulated on HEK cells co-expressing Nrx (blue) and CPTX (green). The right graphs show quantification of GFP/HIS, Nrx (middle) and GluA1–3 (bottom) on HEK cells. The bars represent mean ± SEM. \*\*\**P* < 0.001, n.s., not significant, *n* = 15 fields from 3 independent experiments, one-way ANOVA followed by Tukey’s test. (**B**) Beads coated with recombinant CPTX recruit GluA4 in dendrites of parvalbumin-positive (PV^+^) interneurons. Representative immunocytochemical staining shows that endogenous GluA4 (magenta) in PV^+^ interneurons (green) highly accumulated on contacting beads (arrows) coated with CPTX, but not Cbln1 or vehicle (mock). The right graph shows the quantification of GluA4 immunoreactivities on beads contacting dendrites of PV^+^ interneurons (arrow) subtracted by that on beads non-contacting. The bars represent mean ± SEM values. \*\*\**P* < 0.001, *n* = 25 fields from 4 independent experiments, one-way ANOVA followed by Tukey’s test. (**C**) CPTX, but not Cbln1, colocalize with AMPARs. Representative immunocytochemical staining shows endogenous GluA1–3 (red) and pan-Nrx (green) in hippocampal neurons incubated with vehicle (mock), Cbln1-HIS or CPTX-HIS (blue). Scale bar, 2 μm.

**fig. S6.**
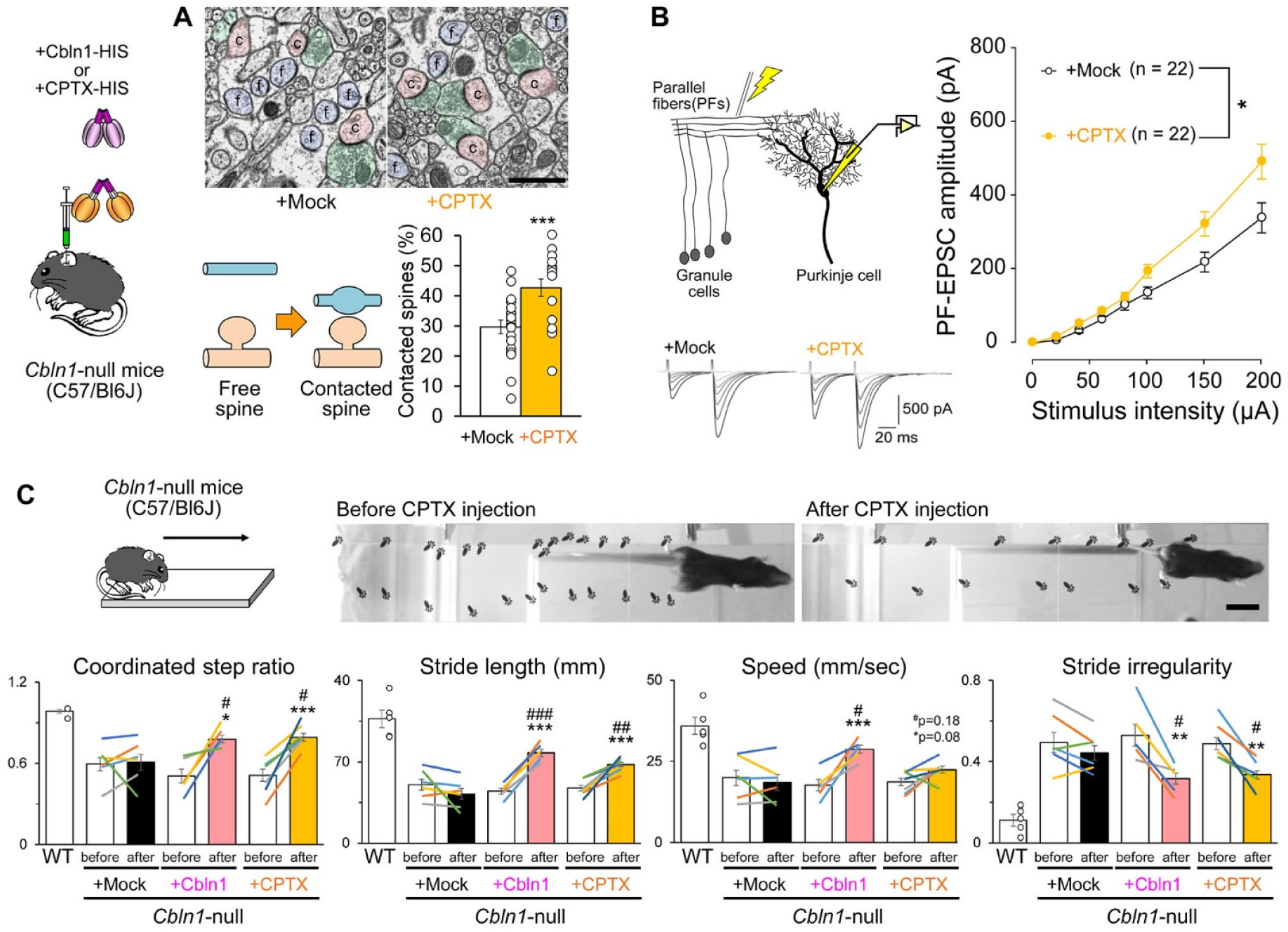
CPTX restores cerebellar PF-PC synapses and improved motor coordination in ***Cbln1-null* mice on the C57BL background**. **(A)** Representative electron microscopic images showing free dendritic spines (light blue) and contacted spines (light red) innervated by PFs (light green) in the cerebellum 3 d after injection of vehicle (mock) or CPTX. Percentages of contacted PC spines are quantified in the lower graph. Scale bar, 500 nm. The bars represent mean ± SEM values. \**P* < 0.01, \*\*\**P* < 0.001, *n* = 19–20 sections from 2 mice, χ^2^ test. **(B)** CPTX restores functional PF-evoked EPSCs in *Cbln1*-null mice 3 d after injection. Representative traces from slices prepared from *Cbln1*-null mice that received vehicle (mock) or CPTX are shown on the top. The graph shows averaged input-output relationship of PF–EPSCs in each treatment. ns, not significant, \**P* < 0.05 *n* = 22 each, two-way ANOVA followed by Scheffe *post-hoc* test. The bars represent mean ± SEM values. **(C)** CPTX improves the gait of *Cbln1*-null mice 3 d after the injection. Representative footprints before and after CPTX injection are shown on the top. Scale bar, 50 mm. The lower graphs show the quantification of gait parameters. The bars are average scores and the lines are an individual score of each mouse before and after vehicle (mock treatment), Cbln1 or CPTX injection. The bars represent mean ± SEM values. \**P* < 0.05, \*\**P* < 0.01, \*\*\**P* < 0.001 compared with before and after by paired *t*-test, ^#^*P* < 0.05, ^##^*P* < 0.01, ^##^*P* < 0.001 compared with after injection of Mock, *n* = 5–7 mice, Student’s *t*-test.

**fig. S7.**
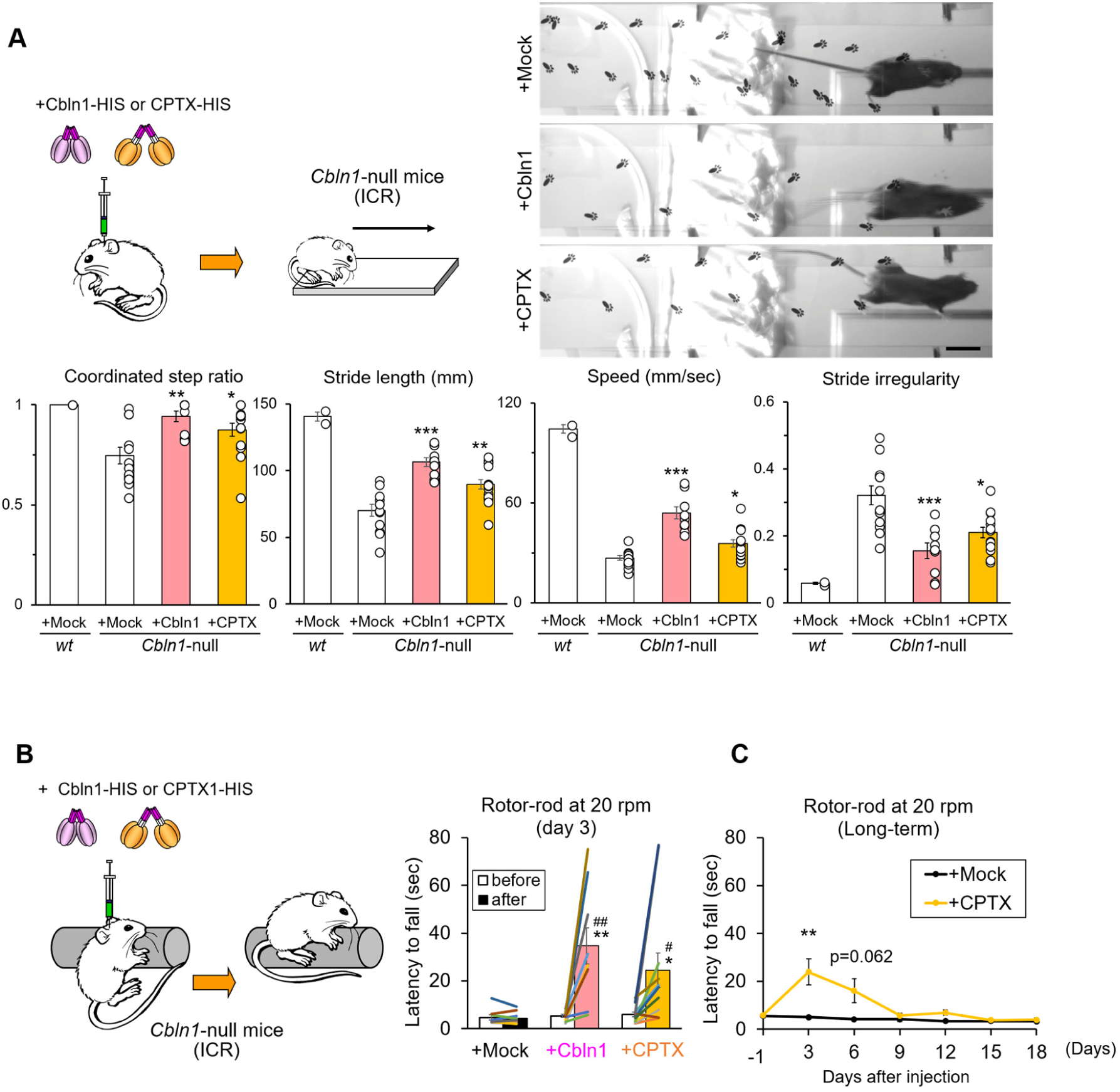
Effects of CPTX on *Cbln1*-null mice on the ICR background. (**A**) Representative footprints of Cbln1-null mice on ICR background on the right. CPTX or Cbln1, but not vehicle (mock treatment) restored gait pattern measured at 3 d after the injection. Scale bar, 50 mm. The lower graphs show the quantification of gait parameters. The bars represent average scores ± SEM. \**P* < 0.05, \*\**P* < 0.01, \*\*\**P* < 0.001 compared with vehicle injection, *n* = 5–7 mice, Student’s *t*-test. (**B**) Cbln1 or CPTX, but not vehicle (mock treatment) restored motor coordination as measured by the rotor-rod test. Latency to fall from a rotating rod (20 rpm) was measured 3 d after the injection. The bars represent the average ± SEM of latency before and after injection. \**P* < 0.05, \*\**P* < 0.01, \*\*\**P* < 0.001 compared with before values by paired t-test, ^#^*P* < 0.05, ^##^*P* < 0.01, compared with mock injections by Student’s t-test. (**C**) The long-term effect of single injections of vehicle (mock treatment) or CPTX on motor coordination. Latency to fall from the rotating rod (20 rpm) was measured every 3 d after the injection. \*\**P* < 0.01 compared with day 0, *n* = 5–21 mice, paired *t*-test.

**fig. S8.**
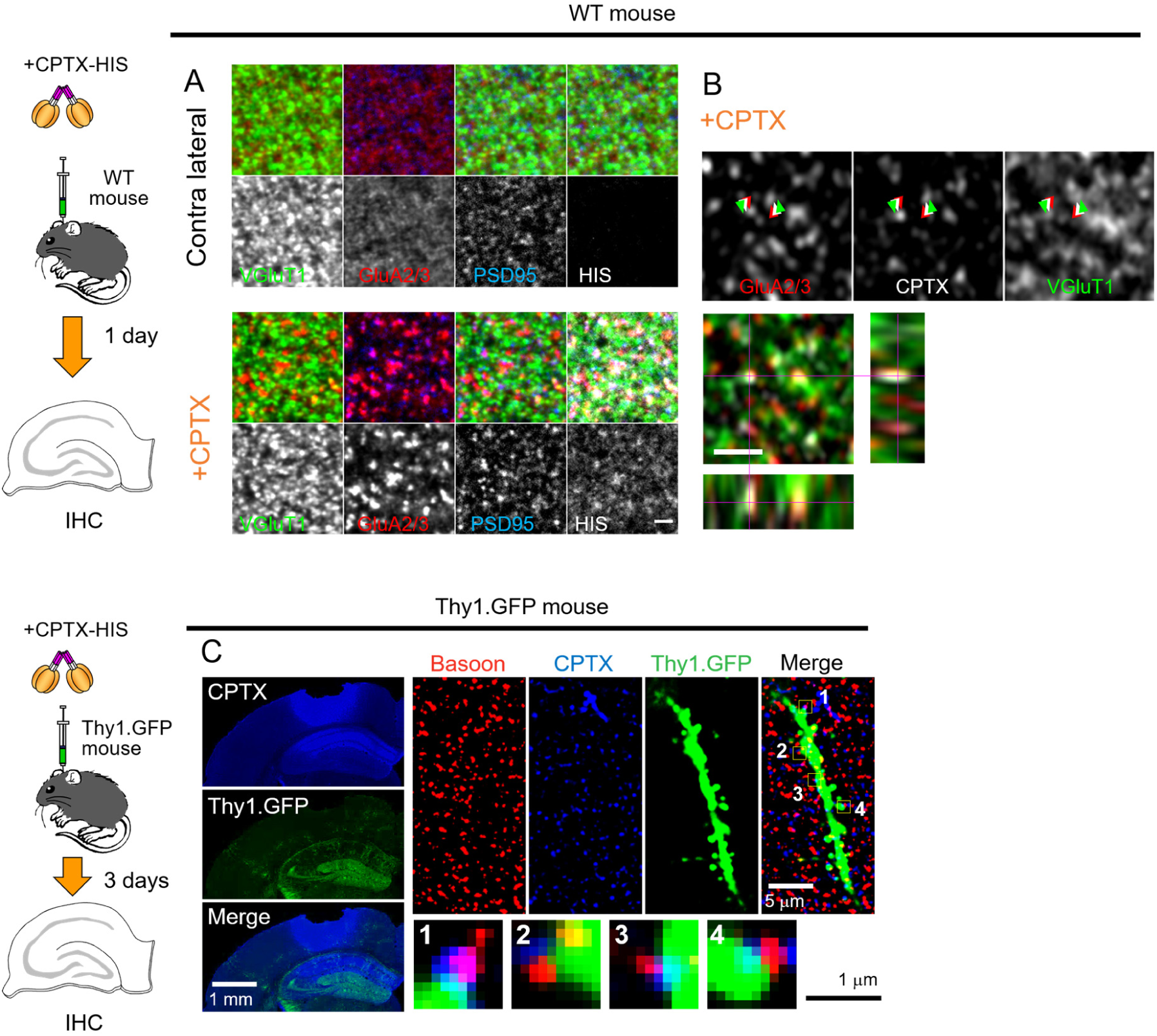
CPTX localizes at excitatory synapses and induces AMPAR accumulation in the hippocampus *in vivo*. (**A**) Representative images at 1 d after injection of CPTX in WT mice. CPTX induced accumulation of GluA2/3 (red) accompanied with PSD95 (blue) and VGluT1 (green) as compared with the contralateral injection site. Scale bar, 2 μm. (**B**) Representative super-resolution microscopic images showing that injected CPTX (gray arrowheads) localizes between VGluT1 (green arrowheads) and GluA2/3 (red arrowheads). Lower panes show reconstituted orthogonal images at red lines. Scale bar, 1 μm. (**C**) Representative images at 3 d after injection of CPTX in Thy1-GFP mice. Left panels show HIS-tagged CPTX (blue) in the CA1 dorsal hippocampus and the cortex. Scale bar, 1 mm. Right panels show magnified immunohistochemical images of CPTX-HIS (blue), bassoon (a presynaptic marker, red) and Thy1-GFP (CA1 neurons). Scale bar, 5 μm. Four boxes indicated in the rightmost panel is enlarged in the bottom panels. Scale bar, 1 μm. CPTX is localized between bassoon-positive presynaptic sites (red) and GFP-positive postsynaptic sites (green) in CA1 neurons.

**fig. S9.**
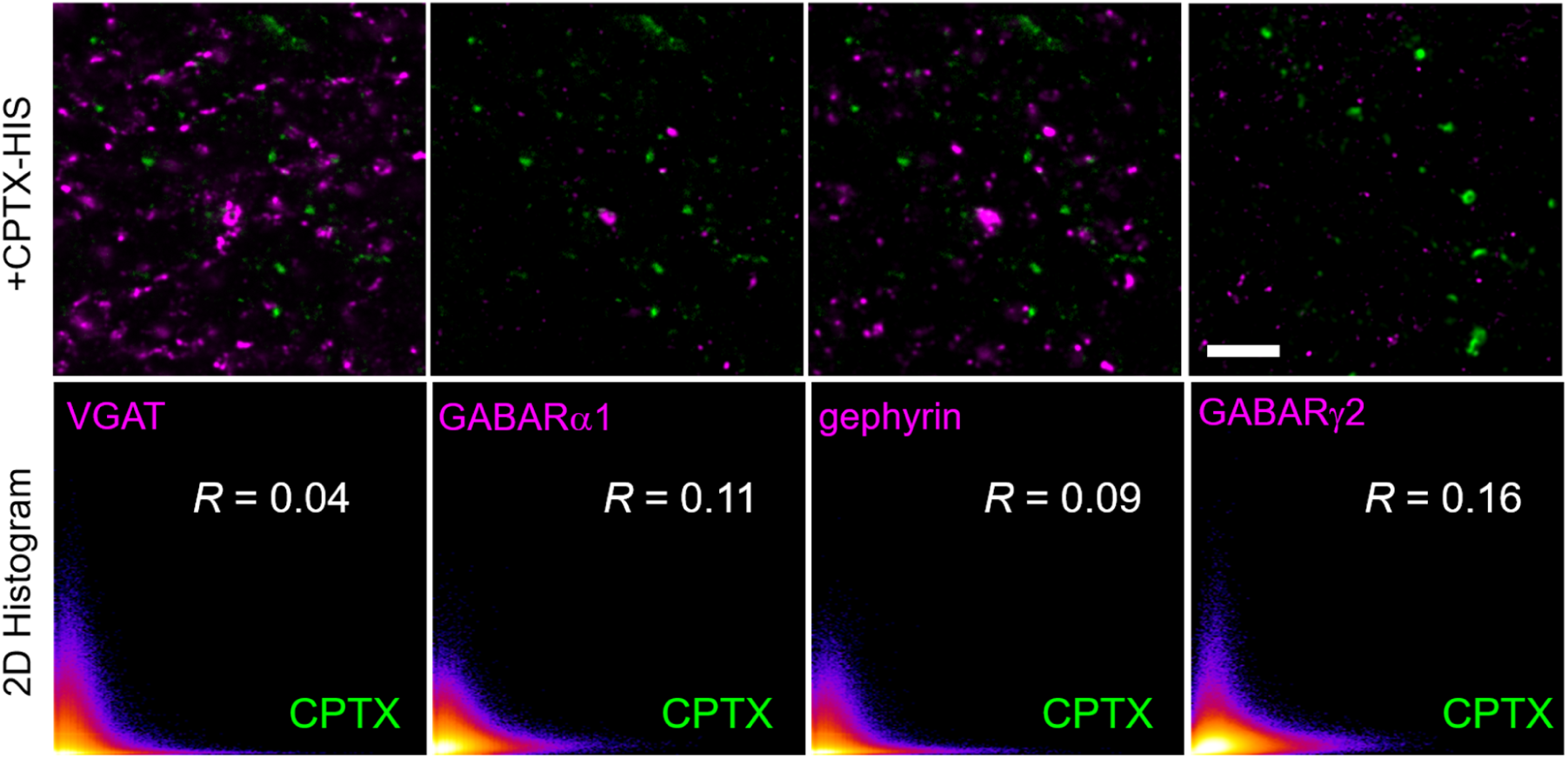
Injected CPTX does not colocalize with inhibitory synapses *in vivo* Representative immunohistochemical images show that CPTX-HIS (green) does not colocalize with inhibitory synapses (magenta) detected by a presynaptic marker VGAT and postsynaptic makers, GABARα1, gephyrin or GABARγ2, at 1 d after the injection of CPTX into the hippocampus. The left three images share the same CPTX signals due to the quadruple staining. Scattered plots show the 2D intensity histogram between CPTX (x-axis) and each inhibitory synapse marker (y-axis) and *R*-value is Pearson’s correlation efficiency. Scale bar, 5 μm.

**fig. S10.**
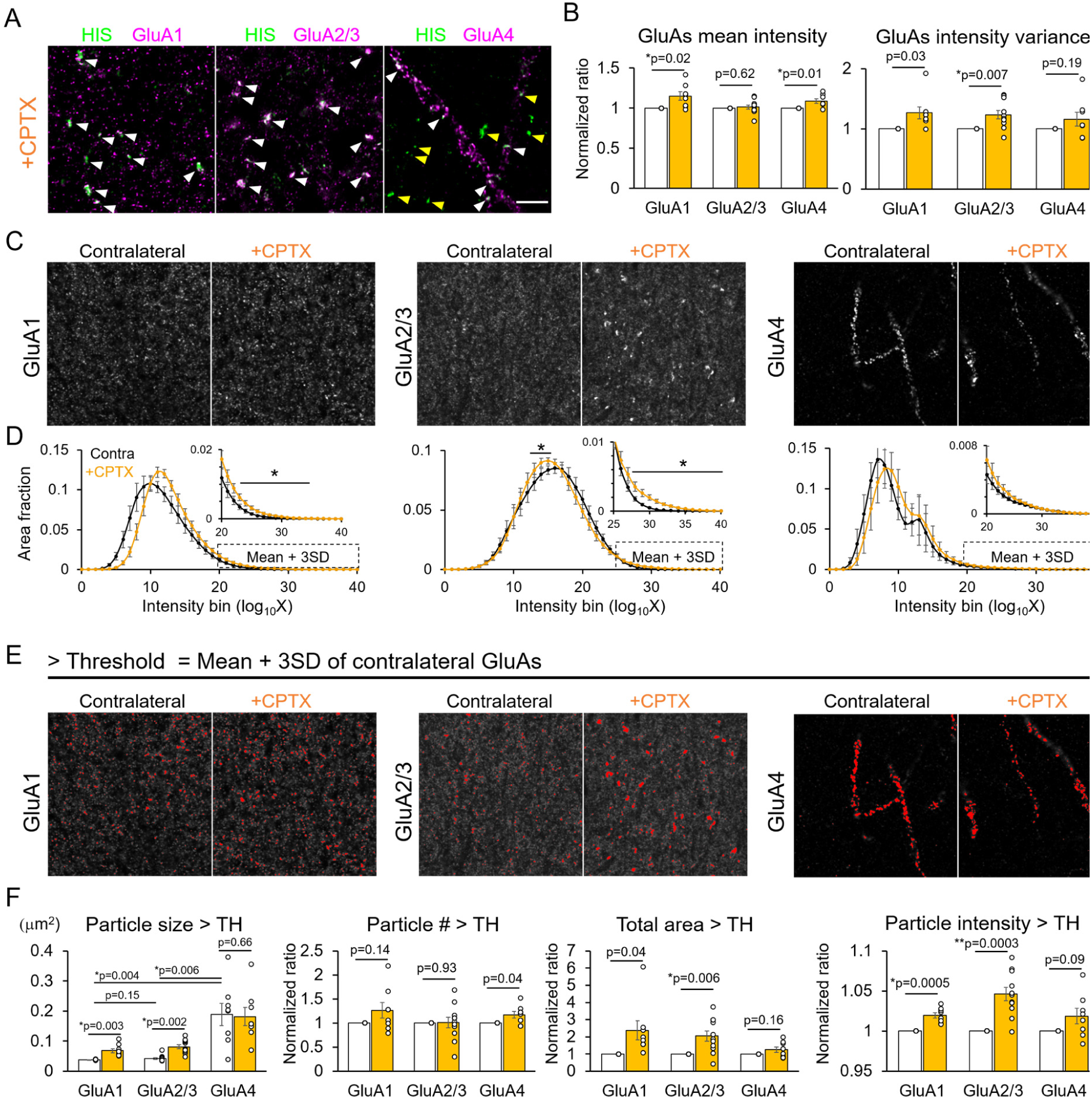
CPTX differentially accumulates GluA subunits in the *stratum radiatum* of the hippocampus *in vivo*. (**A**) Co-immunostaining of HIS (CPTX) and each GluA subunit in the CA1 region of wild-type hippocampus (*stratum radiatum*) 1 d after CPTX injection. White arrowhead indicates the colocalization while yellow arrowhead indicates the CPTX puncta without colocalization with AMPA receptor. Scale bar, 3 μm. (**B**) Quantification of the mean intensity and variance of each GluA subunit. The bar represents the mean ± SEM. \**P* < 0.05, by Student’s *t*-test with Holm correction, *n* = 8–11 images from 3–5 mice. (**C**) Representative GluA1, GluA2/3 and GluA4 immunoreactivities. Contralateral uninjected regions are shown for comparison. Scale bars, 5 μm. (**D**) Normalized intensity histograms of GluA1, GluA2/3 and GluA4 in injected (CPTX) and contralateral (Contra) CA1 regions. While CPTX shifted the histogram of GluA1 and GluA4 to the higher value, it increased the fraction of darker GluA2/3 signals (bins 14–18). Inset shows the magnified area of the histogram corresponding to the bins > mean + 3 SD intensity. \**P* < 0.05, by multi-way analysis of variance (in the bin of mean + 3 SD intensity for GluA1 and GluA4, and in the whole bin for GluA2/3) with post-hoc paired t-test between contra and +CPTX at bins 23–35 in GluA1, bins 13–15 and 27–40 in GluA2/3. Each point represents mean ± SEM. *n* = 8–11 images from 3–5 mice. (**E**) Representative high-intensity immunopositive puncta of each GluA subunit. Pixels that had > mean intensity + 3 SD of the contralateral area are shown in red. (**F**) Quantification of the size, number, total area occupied by particles and intensity of high-intensity (mean + 3 x SD) puncta of each GluA subunit. CPTX increased the size of high-intensity particles of GluA1 and GluA2/3, but not GluA4. Particle number, area and intensity were normalized with those in the contralateral images. \**P* < 0.05, \*\**P* < 0.01, by Student’s *t*-test with Holm correction. The bar represents the mean ± SEM. *n* = 8–11 images from 3–5 mice.

**fig. S11.**
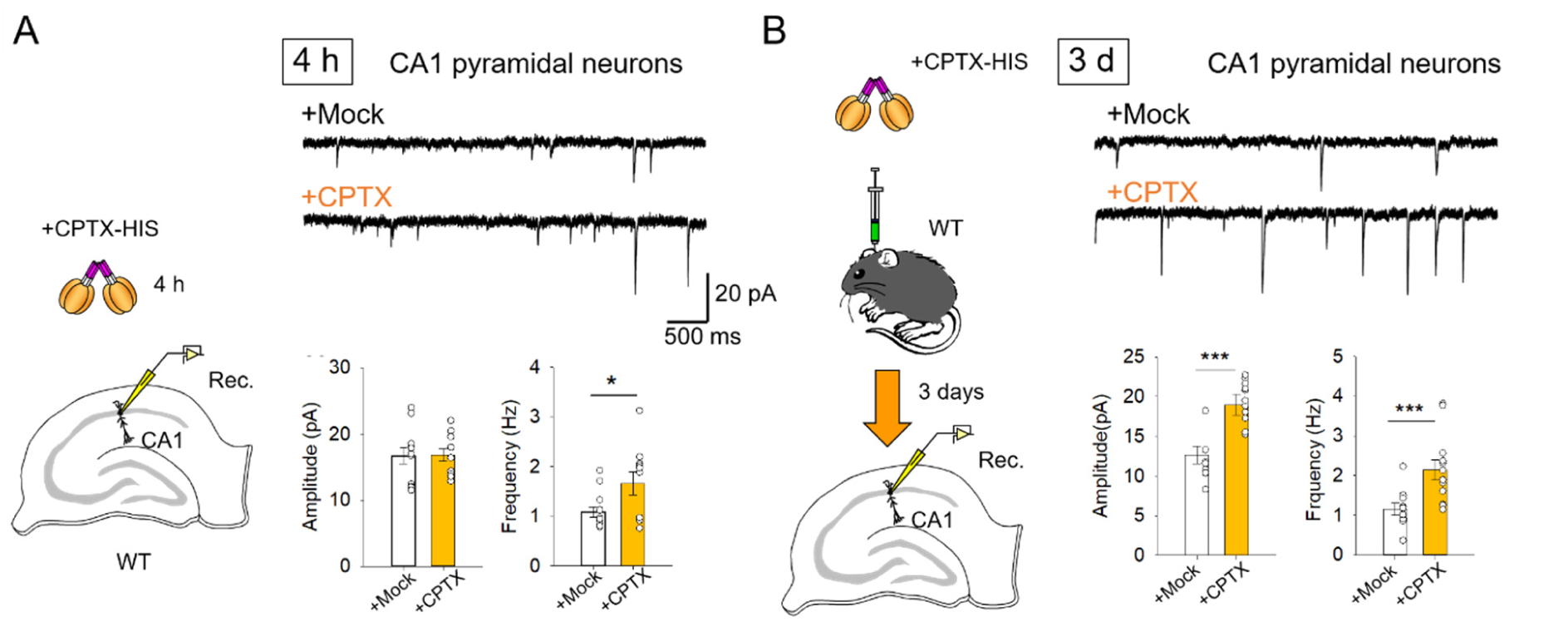
CPTX induces functional excitatory synapses in CA1 hippocampal pyramidal neurons *in vitro* and *in vivo*. **(A)** CPTX treatment for 4 h increases mEPSC frequency in hippocampal slices. Slices acutely prepared from 4- to 6-week-old C57Bl/6J mice were incubated with CPTX (20 μg/mL) or vehicle (mock treatment) for 4 h. Representative traces of mEPSCs are shown on the top. The bottom graphs show the mean + SEM of mEPSC amplitude and frequency. \**P* < 0.05, by Student’s *t*-test. *n* = 12 (Mock (HBS buffer, vehicle of CPTX) and *n* = 11 (CPTX). **(B)** CPTX increases mEPSC amplitude and frequency in hippocampal slices prepared 3d after CPTX injection. Representative traces of mEPSCs are shown on the top. The bottom graphs show the mean + SEM of mEPSC amplitude and frequency. \**P* < 0.05, \*\*\**P* < 0.001, one-way ANOVA followed by Holm-Sidak test, *n* = 10 (Mock) and *n* = 13 (CPTX) from 3 mice each condition.

**fig. S12.**
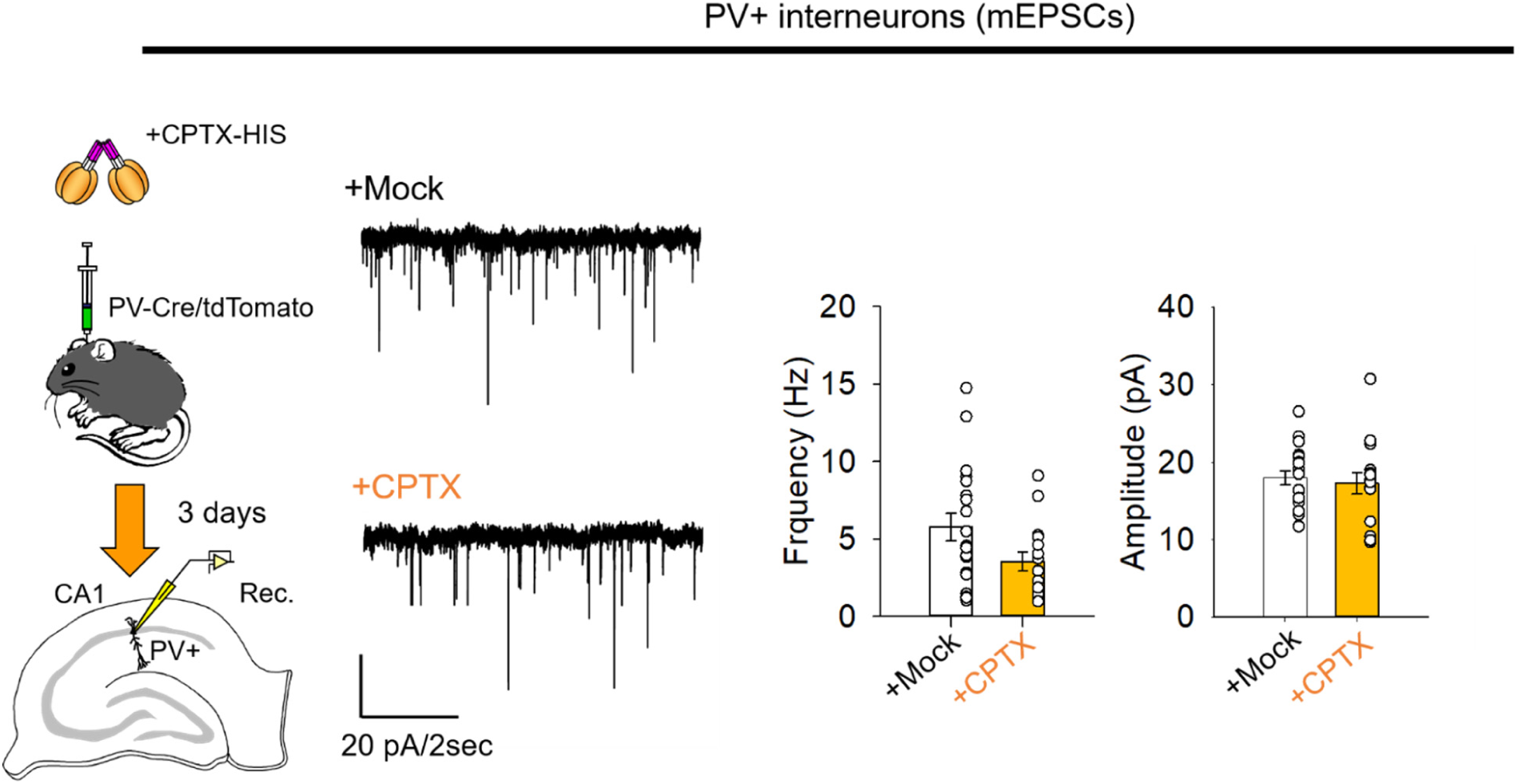
CPTX does not induce excitatory synapses onto PV+ interneurons in the hippocampus *in vivo*. CPTX does not increase mEPSC amplitude and frequency in PV-positive (PV^+^) interneurons in the CA1 region of hippocampal slices prepared 3d after CPTX injection into PV-Cre; TdTomato-LSL transgenic mouse. Representative traces of mEPSCs are shown. The upper graphs show the mean + SEM values of mEPSC amplitude and frequency among different groups. Not statistically significant with Student’s *t*-test, *n* = 19 (Mock (HBS buffer, vehicle of CPTX) and *n* = 15 (CPTX) from 3 mice each.

**fig. S13.**
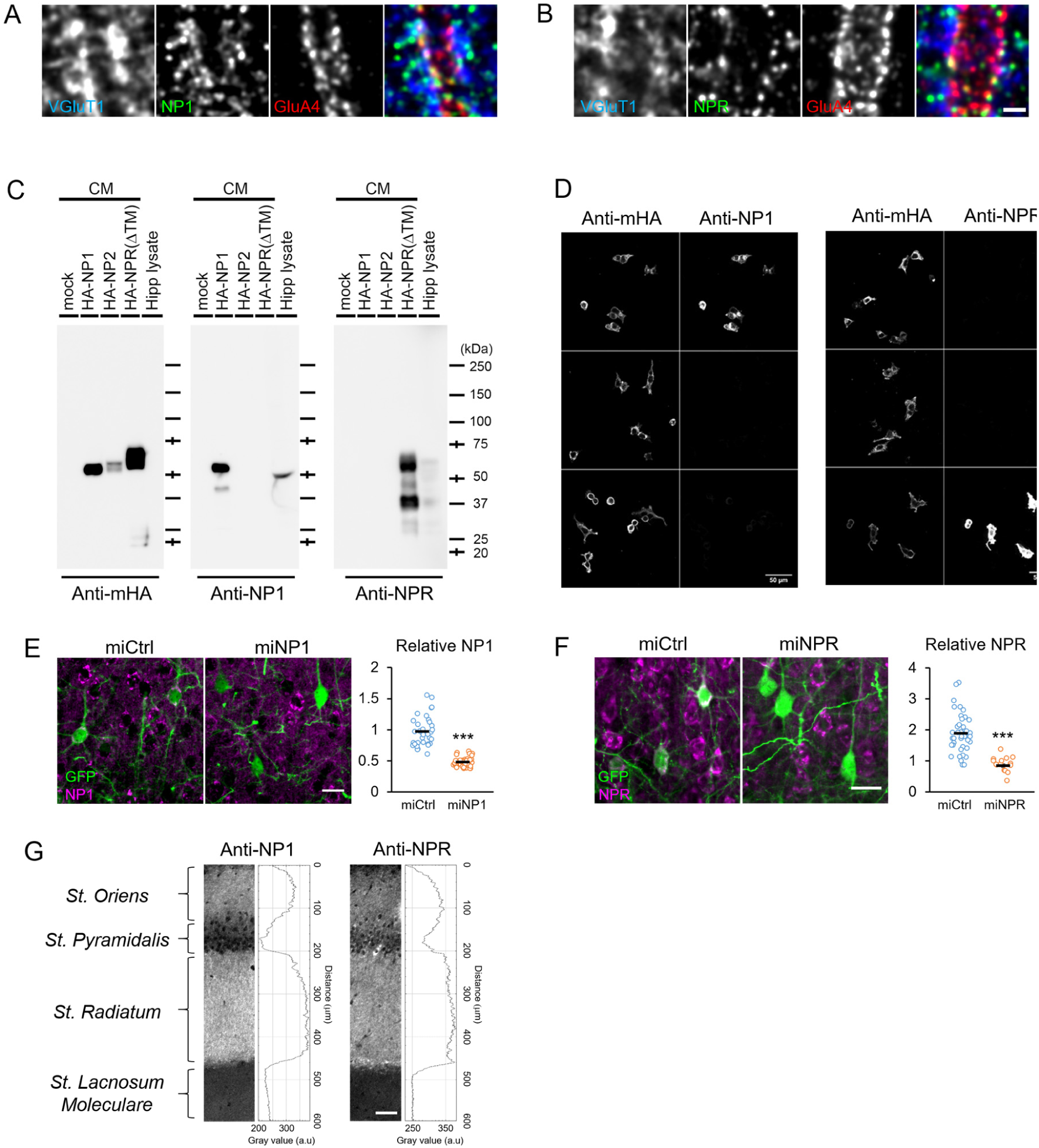
CPTX does not affect excitatory synapses on PV-positive interneurons *in vivo*. (**A, B**) Endogenous NPs highly colocalize with GluA4 in PV^+^ interneurons. Representative immunohistochemical images show endogenous NPs (green; NP1 [A], NPR [B]) colocalize with VGluT1 (blue) and GluA4 (red) in PV^+^ interneurons in the hippocampus. Scale bar, 1 μm (**C–F**). Validation of antibodies against NP1 and NPR. (**C**) Conditioned medium (CM) containing NP1, NP2 or NPR lacking transmembrane domain (ΔTM) expressed in HEK293 cells and hippocampal tissue lysates were analyzed by western blotting with the antibodies against HA, NP1 or NPR. Anti-NP1 and NPR showed the selective band for the overexpressed NP1 or NPR, respectively, and similar size of band in the hippocampal lysate. (**D**) Immunocytochemical analysis of HA-tagged NP1, NP2 and NPR (full-length) expressed in HEK293 cells. Anti-NP1 and NPR antibody showed selective signals for NP1 or NPR, respectively. (**E, F**) Specific immunostaining of endogenous NP1 and NPR in the cortex. cDNAs encoding GFP and control miRNA or miRNAs against NP1 (**E**) or NPR (**F**) were expressed in layer 2/3 cortical neurons by *in utero* electroporation. Representative immunohistochemical images after 4 weeks show a reduction of NPs immunoreactivities (magenta) in neurons (green) expressing miRNA against NPs compared to control miRNAs. Scale bars, 10 μm. The graphs show the quantification of the immunoreactivity of NP1 (**E**) or NPR (**F**). The bars represent mean ± SEM. \*\*\**P* < 0.001, *n* = 15–47 cells, Student’s *t*-test. (**G**) Representative NP1 and NPR immunoreactivities in the hippocampal CA1 region. The intensity histogram is shown for each layer. Scale bar, 50 μm.

**fig. S14.**
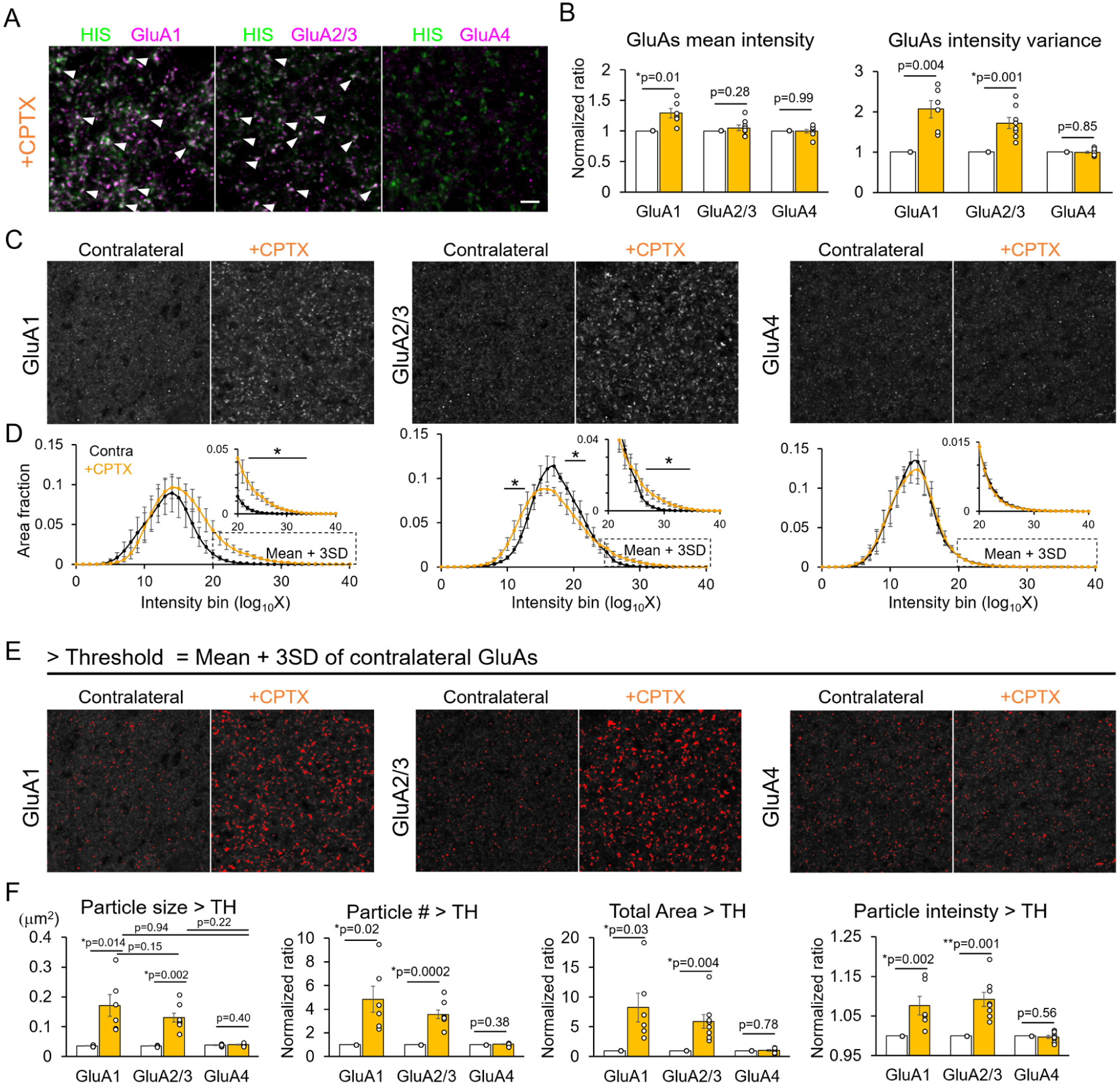
CPTX differentially accumulates GluA subunits in the *stratum lacunosum moleculare* of the hippocampus *in vivo*. (**A**) Co-immunostaining of HIS (CPTX) and each GluA subunit in the CA1 region of wild-type hippocampus (*stratum lacunosum moleculare*) 1 d after CPTX injection. White arrowhead indicates the colocalization of CPTX and GluA1 or GluA2/3. GluA4 signals are weak in this layer. Scale bar, 3 μm. (**B**) Quantification of the mean intensity and variance of each GluA subunit. The bar represents the mean ± SEM. \**P* < 0.05, by Student’s *t*-test with Holm correction, *n* = 8–11 images from 3–5 mice. (**C**) Representative GluA1, GluA2/3 and GluA4 immunoreactivities. Contralateral uninjected regions are shown for comparison. Scale bars, 5 μm. (**D**) Normalized intensity histograms of GluA1, GluA2/3 and GluA4 in injected (CPTX) and contralateral (Contra) CA1 regions. While CPTX shifted the histogram of GluA1 to the higher value, it increased the fraction of darker GluA2/3 signals (bins 14–18). Inset shows the magnified area of the histogram corresponding to the bins > mean + 3 SD intensity. \**P* < 0.05, by multi-way analysis of variance (in the bin of mean + 3 SD intensity for GluA1 and GluA4, and in the whole bin for GluA2/3) with post-hoc paired t-test between contra and +CPTX at bins 22–35 in GluA1, bins 11-13, 18-20 and 25–33 in GluA2/3. Each point represents the mean ± SEM. *n* = 8–11 images from 3–5 mice. The bar represents the mean ± SEM. (**E**) Representative high-intensity immunopositive puncta of each GluA subunit. Pixels that had > mean intensity + 3 SD of the contralateral area are shown in red. (**F**) Quantification of the size, number, total area occupied by particles and intensity of high-intensity (mean + 3 x SD) puncta of each GluA subunit. CPTX increased the size of high-intensity particles of GluA1 and GluA2/3, but not GluA4. Particle number, area and intensity were normalized with those in the contralateral images. \**P* < 0.05, \*\**P* < 0.01, by Student’s *t*-test with Holm correction. The bar represents the mean ± SEM. *n* = 8–11 images from 3–5 mice.

**fig. S15.**
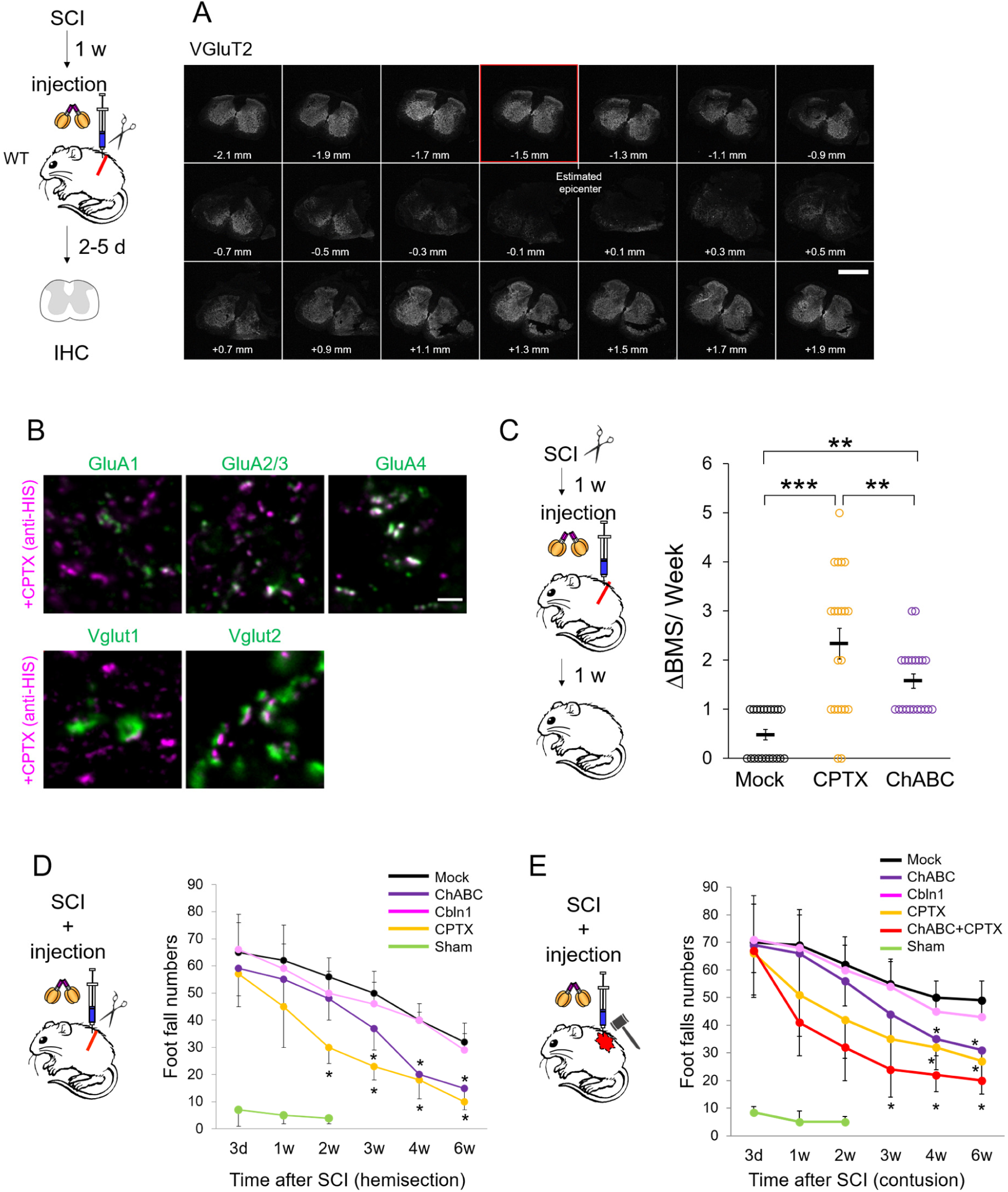
Immunohistochemical and locomotion analyses of spinal cord injury model mice. (**A**) Determination of spared region following SCI. One week after dorsal hemisections, CPTX was injected into the rostral region. Representative immunostaining images of coronal serial sections (20 μm thickness, 200 μm intervals) revealed that sections > 1.1 mm rostral to the epicenter of the injured region showed intact VGluT2 immunoreactivity. Scale bar, 0.5 mm. (**B**) Representative confocal immunohistochemical images of CPTX and synaptic markers. CPTX was mostly colocalized with a postsynaptic marker GluA4 (top) and a presynaptic marker VGluT2 (bottom). Scale bar, 2 μm. (**C**) Rapid recovery of motor performance by CPTX. Mock (HBS buffer, vehicle of CPTX), CPTX or ChABC were delivered 1 week after hemisections and the initial recovery of the motor function in the subsequent week was calculated as ΔBMS/week. The graph shows the mean ± SEM. \*\**P* < 0.01, \*\*\**P* < 0.001, one-way ANOVA followed by Student *t*-test, *n* = 21 mice for each treatment. (**D** and **E**) Time-course of footfall test after hemisection (**C**) and contusion (**D**). Cbln1, CPTX, ChABC or mock (vehicle control) was injected immediately after hemisections. For sham controls, the spinal cord was surgically exposed without imposing injury or injection. The graphs show the mean ± SEM. \**P* < 0.05, Repeated-measures ANOVA followed by Bonferroni-Dunn test, *n* = 6 mice for each treatment.

**Supplemental Table S1.**
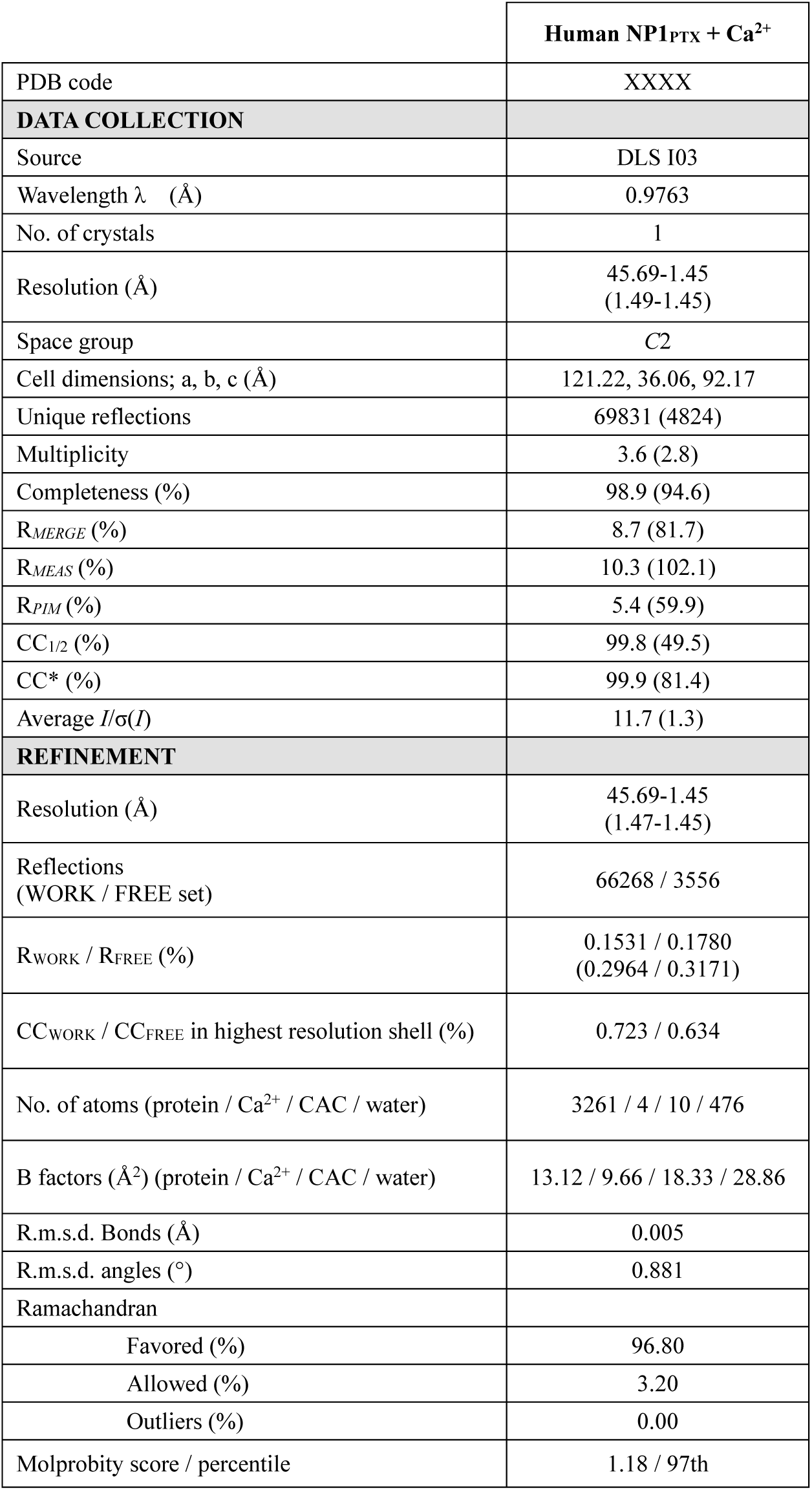
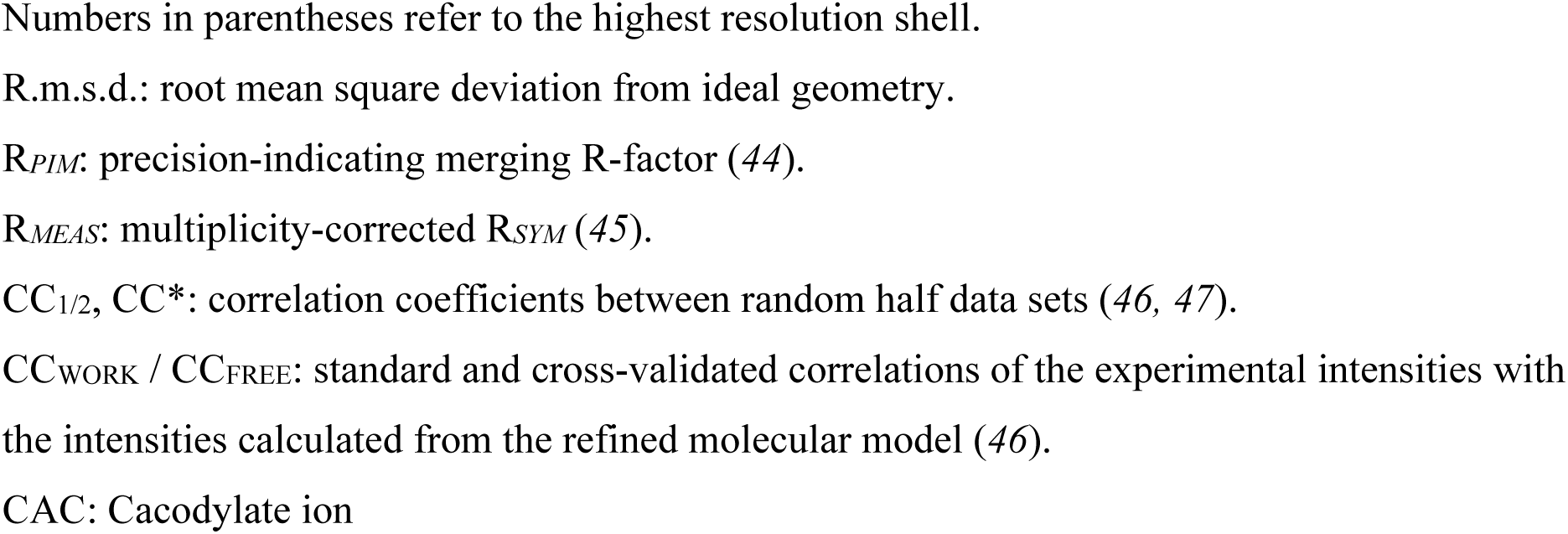
Crystallographic Data Collection and Refinement Statistics.

